# Clinical phenotypes in acute and chronic infarction explained through human ventricular electromechanical modelling and simulations

**DOI:** 10.1101/2022.02.15.480392

**Authors:** Xin Zhou, Zhinuo Jenny Wang, Julia Camps, Jakub Tomek, Alfonso Santiago, Adria Quintanas, Mariano Vazquez, Marmar Vaseghi, Blanca Rodriguez

## Abstract

**Aims:** Sudden death after myocardial infarction (MI) is associated with electrophysiological heterogeneities and ionic remodelling, which are reflected as variable phenotypes. Low ejection fraction (EF) is used in risk stratification, but its mechanistic links with the post-MI pro-arrhythmic heterogeneities are unknown. We aim to provide a mechanistic explanation of clinical phenotypes in acute and chronic MI, from ionic remodeling to ECG and EF, using human electromechanical modelling and simulation to augment experimental and clinical investigations.

**Methods and Results:** A human ventricular electromechanical modelling and simulation framework is constructed and validated with rich experimental and clinical datasets. Abnormalities caused by scar and border zone ionic remodeling are introduced in varying degrees as reported in experimental data obtained in acute and chronic infarction. Simulations enabled reproducing and explaining clinical phenotypes post-MI, from ionic remodelling to ECGs and pressure-volume loops. In acute MI, T-wave inversion and Brugada phenocopy were explained by up to 57 ms of local APD prolongation and activation failure due to the inhibition of potassium, sodium and calcium channels in the border zone. In chronic MI, upright tall T-waves highlight large repolarisation dispersion caused by uneven potassium channel expression in border and remote zones, which promoted ectopic propagation at fast pacing. Post-MI ionic remodelling reduced EF by up to 10% through inhibition of calcium transient amplitude due to weaker calcium currents or SERCA activity, but the EF at resting heart rate was not sensitive to the extent of repolarisation heterogeneity and the risk of repolarisation abnormalities at fast pacing.

**Conclusions:** Multi-scale modelling and simulation coherently integrates experimental and clinical data at subcellular, tissue, and organ scales to unravel electromechanical disease mechanisms in MI. In acute post-MI, ionic remodelling and its effect on refractoriness and propagation failure in the BZ have a strong impact on phenotypic ECG variability, whereas in chronic post-MI, the repolarisation dispersion across the BZ is crucial. T-wave and QT abnormalities are better indicators of repolarisation heterogeneities than EF in post-MI.

## INTRODUCTION

Sudden cardiac death in post-myocardial infarction (post-MI) patients is due to lethal arrhythmias in 50% of cases at both acute and chronic infarct stages (Vazquez *et al*., 2009; Elayi *et al*., 2017). Risk stratification is currently based on low left ventricular ejection fraction (LVEF) (Solomon *et al*., 2005) to identify patients who need the implantation of defibrillator device. However, only a very small subset of patients that suffer from sudden cardiac death are identified by low LVEF, and a significant number of sudden deaths occur in patients with relatively preserved LVEF (Vaduganathan *et al*., 2018). The mechanistic link between electrophysiological heterogeneities, ECG and LVEF phenotypes that underpin arrhythmic events is not clear. A precision medicine approach using computational modelling and simulations could help to elucidate disease mechanisms and provide an *in silico* alternative for therapy evaluation and risk stratification. However, a key hurdle in clinical adoption of such *in silico* tools is providing proof of model credibility through validation studies. In this study, we set out to achieve the dual aim of demonstrating a multi-scale approach for model validation and to identify important electrophysiological mechanisms for post-MI risk stratifications.

Various electrocardiogram (ECG) characteristics were suggested by clinical studies to be relevant to the arrhythmic risk of post-MI patients. Longer QT intervals have been associated with increased mortality as well as ventricular tachycardia and fibrillation for both the acute and chronic MI (Ahnve, 1985; Oikarinen *et al*., 1998). However, QT prolongation in single leads can be a reflection of reduced global repolarisation reserve, while the regional heterogeneity of repolarisation in different post-MI zones can be more crucial (Cluitmans *et al*., 2021) than the global reserve for the development of re-entrant waves. QT dispersion between 12-lead ECGs was proposed as a potential marker for electrophysiological heterogeneity (Spargias *et al*., 1999); however, it was found to have good specificity but low sensitivity (Spargias *et al*., 1999). Prolonged T-peak to T-end interval (Tpe) (Shenthar *et al*., 2015) and increased incidence of T-wave alternans (Bloomfield *et al*., 2004) were also found to be useful risk predictors in post-MI patients. Other ECG metrics, such as QRST integral and spatial ventricular gradients have also shown some promise in improving SCD prediction (Waks *et al*., 2016). However, despite the promising outcomes of those clinical studies, the ECG-based risk predictors are not widely applied in the clinical evaluation of the need for implantable cardioverter-defibrillators (Al-Khatib *et al*., 2018).

A key factor that hinders the clinical utility of ECG biomarkers is the large variability in post-MI phenotypes, their progression from acute to chronic, and between different patients. Variability in QT prolongation (Ahnve, 1985) and post-MI ECG morphologies constrain the use of single biomarker thresholds as predictors. Furthermore, an important limitation of non-invasive ECG biomarkers is their inability to provide direct measurements of regional electrophysiological heterogeneity caused by scar and ionic remodelling, which is critical for the arrhythmic substrate.

Many experimental studies have investigated the underlying mechanisms of increased repolarisation dispersion in post-MI patients. Specifically, repolarisation dispersion increases after MI due to ionic differences between the border zone (BZ) surrounding the scar and the remote zone (RZ) myocardium (Mendonca Costa *et al*., 2018). The BZ at the acute MI exhibits reduced sodium current (I_Na_), L-type calcium current (I_CaL_) (Aggarwal Rajesh & Boyden Penelope A., 1995), and rapid and slow delayed rectifier potassium currents (I_Kr_ and I_Ks_) (Jiang *et al*., 2000), as well as enhanced CaMKII activity (Hund *et al*., 2008a) and gap junction redistribution (Yao *et al*., 2003). After a period of scar healing, ionic remodelling may partially recover (Ursell *et al*., 1985), but increased late sodium current (I_NaL_), calcium-activated potassium and chloride currents (I_KCa_ and I_ClCa_) were observed in the chronic BZ and RZ (Hegyi *et al*., 2018). Post-MI ionic remodelling can cause conduction abnormalities, repolarisation abnormalities, as well as variable alterations in the action potential duration (APD), which act as substrates of ventricular tachycardia (Pogwizd *et al*., 1992). It is, however, unclear how various degrees of repolarisation dispersion affect ECG biomarkers and LVEF used for risk stratification. Therefore, further studies are needed to bridge the gap between cellular electrophysiological characteristics and variable patient phenotypes post-MI. Furthermore, the wealth of cellular, tissue, and ECG data described in the literature provides an excellent multi-scale dataset for model validation. Comparisons of simulated action potential and ECG biomarkers with experimental and clinical data under acute and chronic MI conditions help to improve the credibility of the computational model.

Human-based modelling and simulations of ventricular electrophysiology and mechanics have demonstrated accurate prediction of post-MI arrhythmic risk and myocardial stress and strain (Arevalo *et al*., 2016; Sack *et al*., 2016). However, previous work concentrated on either electrophysiology or mechanics, while the crosstalk between the two, and the relationship between LVEF and arrhythmic risk, is yet to be investigated.

The main goal of this study is, therefore, to quantify the contribution of varying degrees of ionic remodelling to the phenotypic variability in ECG and LVEF biomarkers observed in acute and chronic post-MI patients, and thereby create and validate models of post-MI electromechanics. Using state-of-the-art electromechanical human biventricular simulations, we aim to identify biomarkers that are most representative to the pro-arrhythmic substrate for each state, thus facilitating post-MI risk stratifications to go beyond LVEF. We hypothesise that different ECG abnormalities are explained by different degrees of ionic remodelling leading to activation sequence abnormalities, dispersion of repolarisation, early after depolarisations (EADs) and alternans, whereas LVEF is insensitive to ionic remodelling that underpins ECG disease markers and reflects predominantly calcium transient and structural abnormalities.

## METHODS

### 1. Human multi-scale ventricular electromechanical modelling and simulation: from ionic remodelling to ECG and LVEF

A human ventricular electromechanical modelling and simulation framework is constructed using a population of models approach and evaluated using experimental and clinical data to enable the investigations of variable post-MI patient phenotypes, from ionic remodelling to body surface ECGs and pressure-volume (PV) loops (Figure 1A). A cardiac magnetic resonance (CMR)-based biventricular anatomical mesh (Figure 1B) with corresponding torso geometry was used for all simulations in this study, with an anterior scar that is 75% transmural (Wang *et al*., 2021). Electrical propagation was simulated using the monodomain equation with orthotropic diffusion based on rule-based fields for fibre directions (Streeter *et al*., 1970) with sheet directions normal to the endocardial/epicardial surface (Levrero-Florencio *et al*., 2020) (Figure 1B). Transmural and apex-to-base heterogeneities (Mincholé *et al*., 2019) were introduced and a sensitivity analysis was performed to investigate their implications in ECG biomarkers and LVEF (Figure 1B). Electrical stimulus via Purkinje-myocardial junctions was simulated by an endocardial fast-activation layer with root node locations to achieve realistic QRS complex morphologies simulated at clinically standard lead locations (Figure 1C) (Mincholé *et al*., 2019). In healthy tissue (normal zone (NZ)), monodomain diffusivities were calibrated to achieve experimentally measured orthotropic conduction velocities of 67 cm/s, 30 cm/s, and 17 cm/s (Caldwell *et al*., 2009).

**Figure 1:**
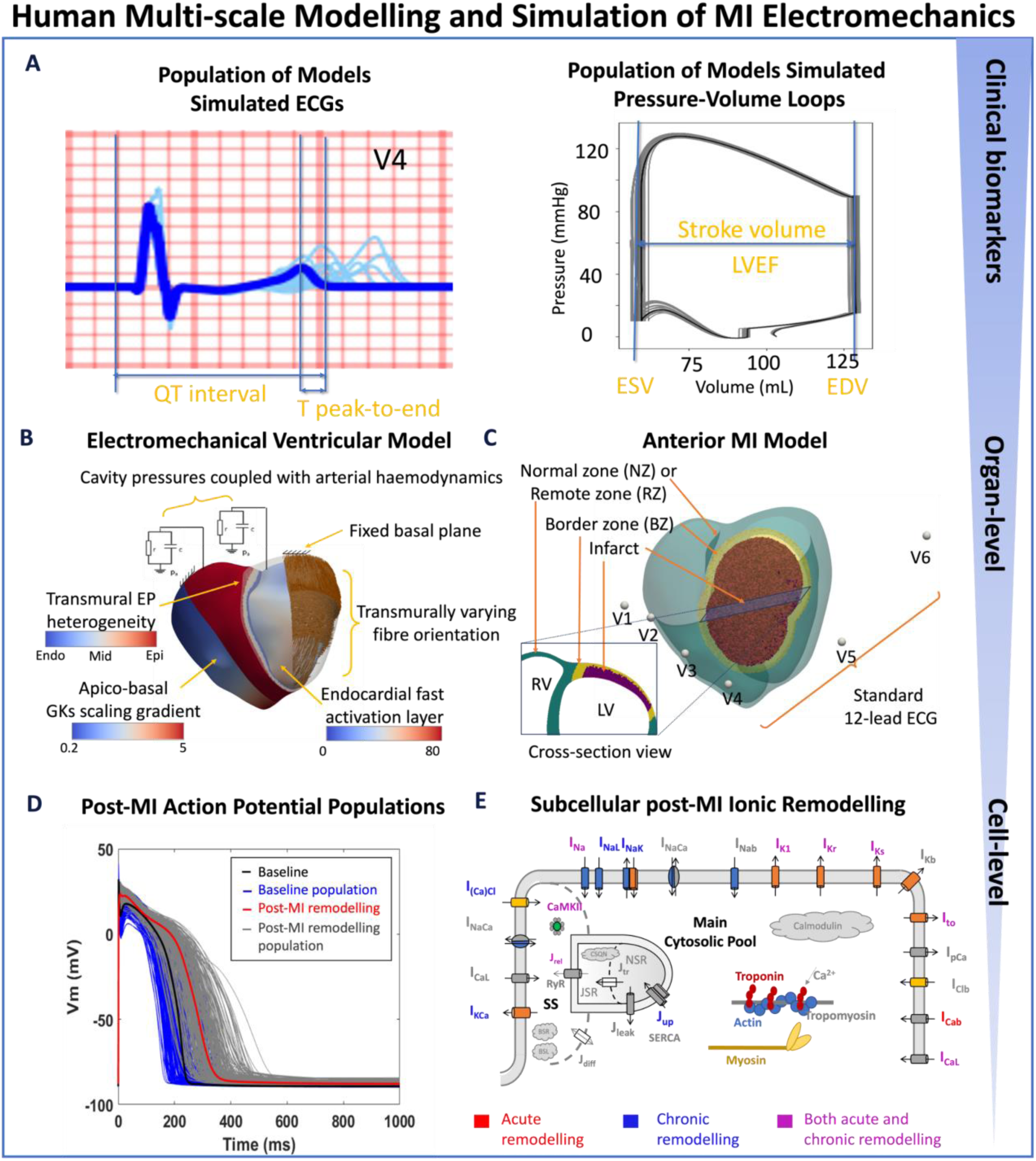
Human-based multi-scale modelling and simulation in acute and chronic myocardial infarction. (A) Simulations using a population of ventricular models (n=17) to produce ECGs (light blue traces) and pressure-volume (PV) loops (grey traces) superimposed with the baseline ventricular model (ECG in blue and PV in black). Biomarkers are calculated from the baseline simulation of ECG and PV, as illustrated. (B) Ventricular electrophysiology is simulated using a fast endocardial activation layer to approximate Purkinje-myocardial junction, experimentally-informed transmural and apico-basal heterogeneities in action potential duration, and transmurally varying myocyte orientation. Mechanical pumping behaviour is modelled by coupling the intraventricular pressures with a two-element Windkessel model of arterial haemodynamics with a fixed basal plane. (C) An anterior 75% transmural infarction is modelled with acute and chronic ionic remodelling embedded in the border zone and remote zones. Standard 12-lead ECG was evaluated at standard body-surface locations. (D) Simulated action potentials using populations of human ventricular models in healthy (baseline) and acute and chronic post-MI conditions with different degrees of ionic remodelling. (E) Schematic representation of ionic fluxes, calcium dynamics and actin/myosin contraction mechanisms in the human ventricular electromechanically-coupled cellular model.

Since anterior infarction is very common and related to the worst prognosis (Stone *et al*., 1988), 75% transmural extent of infarct from the endocardial surface was introduced to the anterior myocardial wall (Figure 1C) to match the clinical definition of transmural infarctions (Sato *et al*., 2008), with scar and border zones (maximum width 0.5 cm) taking up 12.4% and 11.7% of ventricular volume (or 15.8% and 15.0% of left ventricular volume), respectively. Maximum border zone width was based on various reports of systolic strain dysfunction in < 1 cm proximity to the infarct (Gallagher *et al*., 1986; Van Leuven *et al*., 1994). Electrophysiological remodelling was implemented in the border zone (BZ) and the chronic remote zone (RZ), which covered the entire non-infarcted and non-BZ region in both ventricles. In BZ and infarct zones, the diffusivities were calibrated to reproduce conduction slowing (one-third of the NZ conduction velocities). For each virtual cardiomyocyte, the electromechanical single cell model considered the human-based ToR-ORd electrophysiological model (Tomek *et al*., 2019) (extensively validated in control, disease and drug block conditions) coupled with human excitation-contraction and active tension Land model (Land *et al*., 2017; Levrero-Florencio *et al*., 2020) (Figure 1E).

The human biventricular model incorporated strongly-coupled electromechanics with orthotropic passive mechanical behaviour and balance of linear momentum with inertial effects, as in our previous work (Levrero-Florencio *et al*., 2020; Margara *et al*., 2021; Wang *et al*., 2021). In brief, firstly, intracellular calcium concentration drives crossbridge cycling and force production through unblocking the crossbridge binding-site. Secondly, calcium sensitivity and force production are a function of fibre stretch ratio. Thirdly, stretch rate affects distortion-dependent cross-bridge unbinding (Land *et al*., 2017) and the diffusivity tensor is transformed using the deformation gradient tensor such that the prescribed values describe conductivities in the deformed state (Levrero-Florencio *et al*., 2020).

An elastic spring boundary condition was set to act perpendicularly to the epicardial surface, to simulate pericardial constraint, and the basal plane was fixed in space to prevent unphysiological tilting and expansion due to the unavailability of closed basal geometry. Pressure boundary condition on the left and right endocardial surfaces were controlled using a series of five equations (see SM2) that controls 1) active diastolic inflation followed by electrical activation and 2) isovolumic contraction, 3) ejection coupled with two-element Windkessel aortic haemodynamics model, 4) isovolumic relaxation, and 5) passive inflation and relaxation (Wang *et al*., 2021).

The sheet active tension was set to 30% of fibre active tension to achieve sufficient LVEF in control conditions, based on the results from a sensitivity analysis in Supplementary Material (SM) section SM5. In brief, we evaluated the sensitivity of ECG morphology and LVEF to changes in the calcium sensitivity of troponin binding in excitation-contraction coupling and the active tension in the sheet direction as a percentage of that in the fibre direction.

The passive stiffness parameters were calibrated based on a previous sensitivity analysis (Wang *et al*., 2021) to achieve 53% LVEF and a physiological pressure-volume (PV) loop in a control simulation (see SM Table S2 and S3 for a list of calibrated parameters).

The active tension was set to zero in the scar to represent the myocyte damage, and the chronic passive stiffness parameters of the infarct region were increased 10-fold to mimic fibrotic scar formation (Sun *et al*., 2009). For each acute post-MI phenotype, a case of complete loss of contractile function (zero active tension) in the BZ was also simulated to evaluate the contribution of other non-ionic remodelling-related abnormalities on ejection fraction.

### 2. Experimentally-informed single cell and ventricular populations of human post-MI electromechanical models

To account for the inter-subject electrophysiological variability widely observed in clinical data, the baseline human cellular electromechanical ToR-Land model was extended to populations of healthy cellular models. Then, post-MI ionic remodelling was applied to generate populations of post-MI virtual cardiomyocytes (Figure 1D). In addition to the baseline ToR-ORd model, several representative cellular models were selected from the population and implemented into the biventricular electromechanical simulations.

An initial population of human ventricular cell models was constructed based on the ToR-ORd model by varying the conductances or magnitudes of I_Na_, I_NaL_, I_to_, I_CaL_, I_Kr_, I_Ks,_ I_K1_, I_NaCa_, I_NaK_, J_rel_ and J_up_ by up to ±50% using Latin Hypercube Sampling (SM Figure S1 and S2). As illustrated in previous studies, small populations of models with proarrhythmic ionic remodelling achieved similar predictability as large populations with uniform current variations (Zhou *et al*., 2019). Therefore, we chose to start with a small healthy population (n=500) and then introduce multiple combinations of ionic remodelling to mimic the large post-MI variability. After calibration with human experimental data (SM Table S1) and discarding those manifesting EADs at 1Hz, 245 sets of endocardial, midmyocardial and epicardial models were accepted as the healthy population (SM Figure S2). From this population, a total of 17 sets of cell models were randomly selected and uniformly embedded in 17 ventricular models, to generate ventricular population of models that produce a variety of ECGs (Figure 1A, light blue traces) and PV loops (Figure 1B, grey traces).

Several degrees of post-MI ionic remodelling were collated from a combination of human and animal experimental data with variability in severity of disease to explore whether such variabilities can explain variability in established clinical ECG phenotypes. These remodellings have been applied to the healthy celllular model population (n=245) to generate BZ and RZ populations for both acute and chronic post-MI. For acute post-MI (within a week post-occlusion), three types of BZ remodelling (Acute BZ1-3) were considered based on previous modelling work and experimental canine data collected within 5 days post-MI. The three models of acute border zone remodelling had in common strong inhibition of I_Na_ (60∼62%). The BZ2 model had more severe inhibition of I_CaL_ and I_Kr_ than the BZ1 model, alongside other minor differences. The BZ3 model, while having less severe potassium currents inhibition than BZ1, had additional remodellings in CaMKII dynamics, RyR time constants, and I_Cab_. For chronic post-MI, ionic remodelling measured from minipigs 5 months post-MI with heart failure were used to generate Chronic BZ (affecting only the BZ) and Chronic RZ1 (also affecting the remote myocardium). Another type of remodelling, Chronic RZ2, was established based on multiple experimental data from failing human cardiomyocytes. The RZ covers the entire myocardium apart from the infarct and BZ in chronic MI simulations. Furthermore, reduction of sodium current and SERCA, with enhanced CaMKII activity and slower calcium release induced by CaMKII activation were also implemented in Chronic BZ, RZ1 and RZ2, as observed in human failing cardiomyocytes (details in SM Table S4). Compared with RZ1, the RZ2 model had a significantly stronger inhibition of potassium currents and a lower repolarisation reserve, alongside other more minor differences. Both human recordings and animal data were used for model evaluation, given the scarcity of human tissue, summarised in SM Table S4-S5. Action potential, calcium transient, and active tension characteristics of the post-MI models are summarised in SM Table S6. These post-MI ionic remodellings were then applied to the ventricular population of models (n=17) to explain ECG and PV phenotypes while considering physiological population variability in baseline ionic conductances. These remodelled cell models are embedded uniformly within each region according to a prescribed transmural heterogeneity of 30% endo, 40% mid-myocardial, and 30% epicardial cell types.

### 3. Simulation protocols and biomarker calculation

Human virtual ventricular myocytes were paced at 1Hz for 500 beats to detect EAD generation. For alternans generation, single cells were paced at cycle lengths (CLs) of 500 ms, 400 ms and 300 ms for 500 beats, and a ΔAPD greater than 3 ms between the last two beats at steady state was defined as alternans.

Biventricular electromechanical simulations were performed at 800 ms CL (75 beats per minute) with 100 ms allowed for active diastolic filling prior to endocardial activation for each beat. Three beats were sufficient to achieve converged ECG and PV characteristics at 800 ms CL (see Figure S7 for ECGs for all beats).

For chronic post-MI, an additional fast pacing protocol was applied with 500 ms CL (120 beats per minute), with 50 ms allowed for active diastolic filling for each beat. From these fast pacing chronic MI simulations, two sets of cell models showing EAD and alternans behaviours, respectively, were selected from the population of models and embedded according to transmural heterogeneity in the remote zone for ventricular simulations, to test whether the arrhythmic behaviours at the cellular level can result in arrhythmic behaviour at ventricular level and manifest in the ECG. For this fast pacing protocol, six beats were necessary to achieve converged ECG and PV characteristics at 500 ms CL (see Figure S8 for simulated ECGs of all beats).

Clinical biomarkers were quantified from the simulated ECG (including the QT interval, QRS duration, T-wave duration, T peak to T end duration, T onset to T peak duration, and QT dispersion, see definition and method of evaluation in SM2 and biomarker results in SM6 Table S8), and from the simulated PV loop (including end diastolic and end systolic volumes, peak systolic pressures and LV and RV ejection fractions), as well as wall thickening strain (see Table S8-S9 for ECG and PV biomarkers for all simulated beats).

### 4. Simulation software and computational framework

Cellular electrophysiological simulations and Latin Hypercube Sampling were performed using bespoke MATLAB codes. Coupled cellular electromechanics, as well as biventricular electromechanics simulations, were performed using the high-performance numerical software, Alya, for complex coupled multi-physics and multi-scale problems (Santiago *et al*., 2018) on the CSCS (Swiss National Supercomputing Centre) Piz Daint supercomputer multi-core clusters, granted through the PRACE (Partnership for Advanced Computing in Europe) project. The simulation input files and Alya executable required to replicate the simulated results are available upon request for scientific investigations.

### 5. Data availability

The simulation input files and Alya executable required to replicate the simulated results will be made available upon request. Python and MATLAB scripts for cellular populations of models simulations and post-processing of the biventricular simulations can be found at: https://github.com/jennyhelyanwe/post_MI_postprocessing

## RESULTS

### 1. Human modelling and simulation for ECG phenotypes in acute and chronic post-MI

Figure 2 demonstrates the ability of human electromechanical simulations to reproduce a variety of clinically reported phenotypes in patients with acute and chronic infarction, in agreement with clinical measurements of ECG and pressure-volume biomarkers, as quantified in further detail in (SM Table S7, details of clinical database in SM4). When imposing acute post-MI remodelling, simulated ECGs reproduced fractionated QRS complexes, T-wave inversion, Brugada phenocopy ST-segment elevation and QT interval prolongation in the anterior leads (Figure 2A), which are common ECG phenotypes observed in acute post-MI patients. Simulations also recapitulated ECG morphology similar to healthy subjects with upright T-waves (Figure 2A, right), which can also be present in acute post-MI.

**Figure 2:**
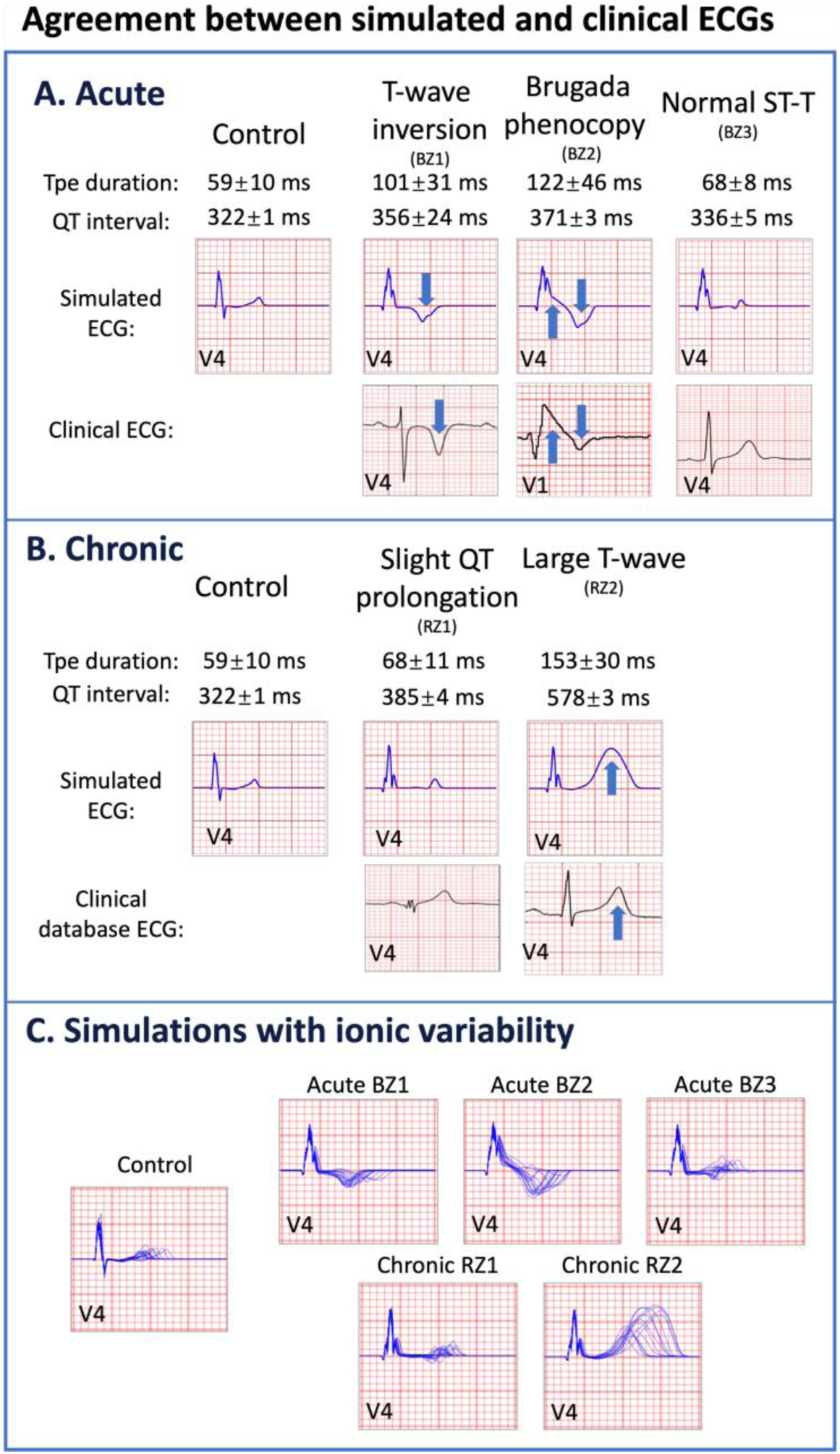
Agreement between simulated and clinical ECGs demonstrating variability in clinical phenotypes in acute and chronic post-myocardial infarction (post-MI). (A) In acute MI, simulated ECGs show T-wave inversion (border zone model 1 (BZ1)), Brugada phenocopy (BZ2), and normal phenotypes (BZ3), in accordance with phenotypes found in clinical databases. (B) In chronic MI, simulated ECGs show prolonged QT and upright T-waves with a range of amplitude and duration (remote zone model 1 and 2 (RZ1, RZ2)) comparable to those observed in clinical databases. (C) ECG simulations of control, and acute and chronic post-MI considering ionic variability of the baseline ToR-ORd model. T wave morphologies for acute and chronic post-MI are mostly preserved across ionic variability.

In chronic post-MI, simulated ECG displayed upright T-waves, and global prolongations of QT intervals in all precordial leads (Figure 2B, top row), which are characteristic of chronic patients with worse clinical outcomes (Ahnve, 1985; Oikarinen *et al*., 1998). A range of T-wave durations were present, as can be found in chronic post-MI (Figure 2B, bottom row). SM4 summarises the clinical ECGs used for comparisons in Figure 2. The full 12-lead ECGs for acute and chronic post-MI simulations can be found in SM Figure S7 and S8.

Simulations using the ventricular population of models showed that the described ECG features of acute and chronic post-MI were mostly preserved across variations in ionic conductances (Figure 2C). Sensitivity analysis showed first that large changes in apex-to-base and transmural heterogeneities only altered T-wave amplitude but not its polarity and did not affect the ST-segment (SM Figure S3 and S4), and second that changes in mechanical parameters did not affect the ECG morphology (SM Figure S5 and S6). This result supports the specificity of post-MI signatures to underlying ionic remodelling.

### 2. In acute MI, T-wave inversion and Brugada phenocopy can indicate reversed transmural repolarisation gradient and activation failure

Analysis of simulation results enabled the uncovering of specific contributions of different degrees of ionic remodelling to ECG phenotypes identified in acute versus chronic MI (Figure 3 vs Figure 4). In acute MI, T-wave inversion phenotype was associated with a reversed transmural repolarisation gradient (Figure 3A, repolarisation time map insets), due to 57 ms of APD prolongation at the epicardial BZ compared with control (Figure 3B, membrane potential). APD prolongation originated due to inhibition of multiple potassium currents caused by BZ1 ionic remodellling (Figure 3C, the first column of I_Kr_, and SM Table S4 Acute BZ1).

**Figure 3:**
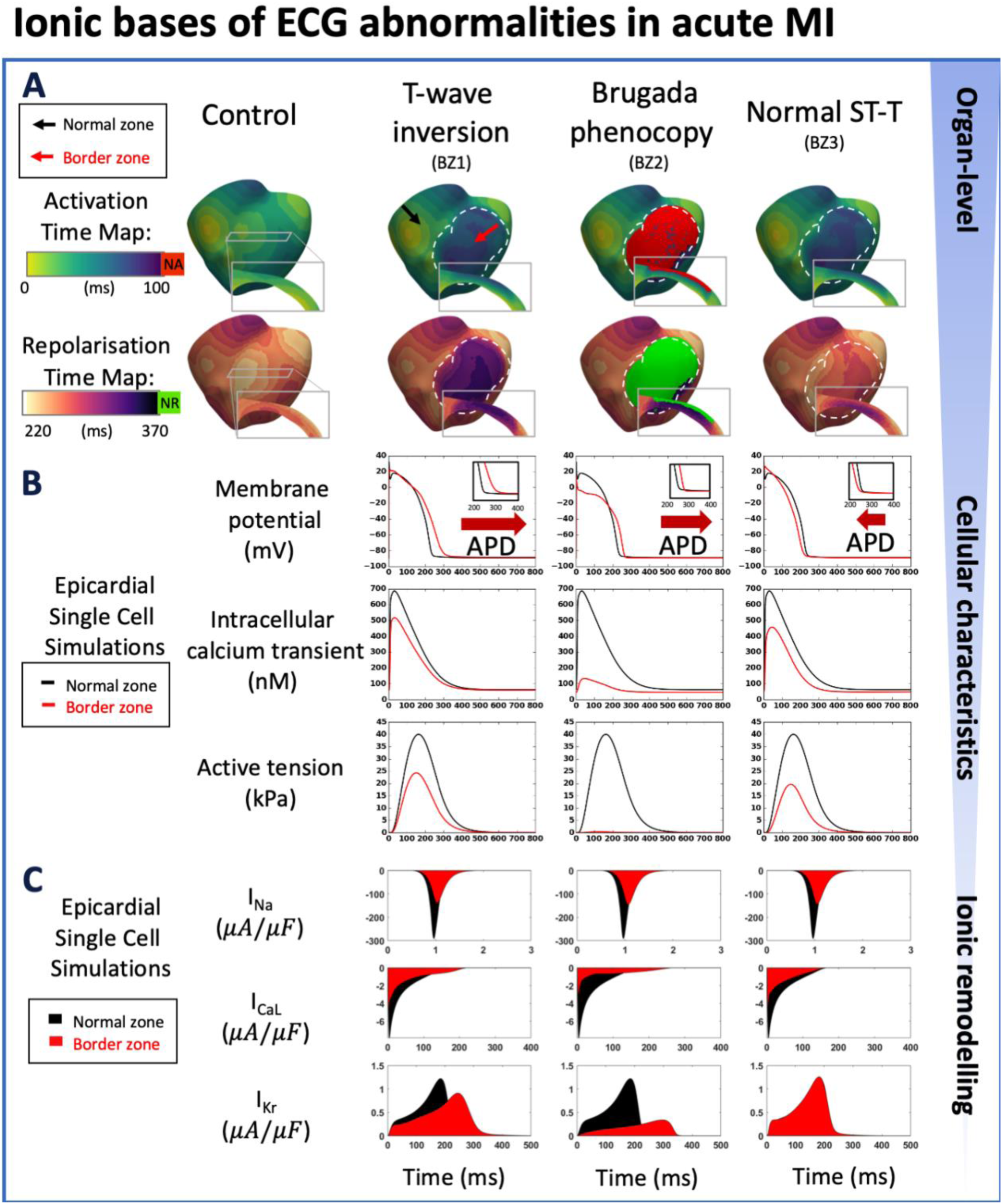
Multiscale explanation of ST and T-wave phenotypes in acute MI. (A) Activation time maps reveal conduction delay in acute border zone in T-wave inversion and normal ST-T phenotypes, and conduction block in Brugada phenocopy, as well as large repolarisation dispersion and altered transmural repolarisation gradient in T-wave inversion and Brugada phenocopy. Red in activation map show regions of no activation (NA), green in repolarisation map highlights regions of no repolarisation (NR). (B) Action potential duration (APD) prolongation is present in T-wave inversion and Brugada phenocopy cellular phenotypes, as well as slight shortening (normal ST-T) (red arrows), with decreased calcium amplitude in all phenotypes, and a corresponding decrease in active tension generation. (C) I_Na_, I_CaL_, and I_Kr_ remodelling underpin reduced conduction, reduced calcium amplitude, and alterations in action potential duration, respectively, in all acute phenotypes.

**Figure 4:**
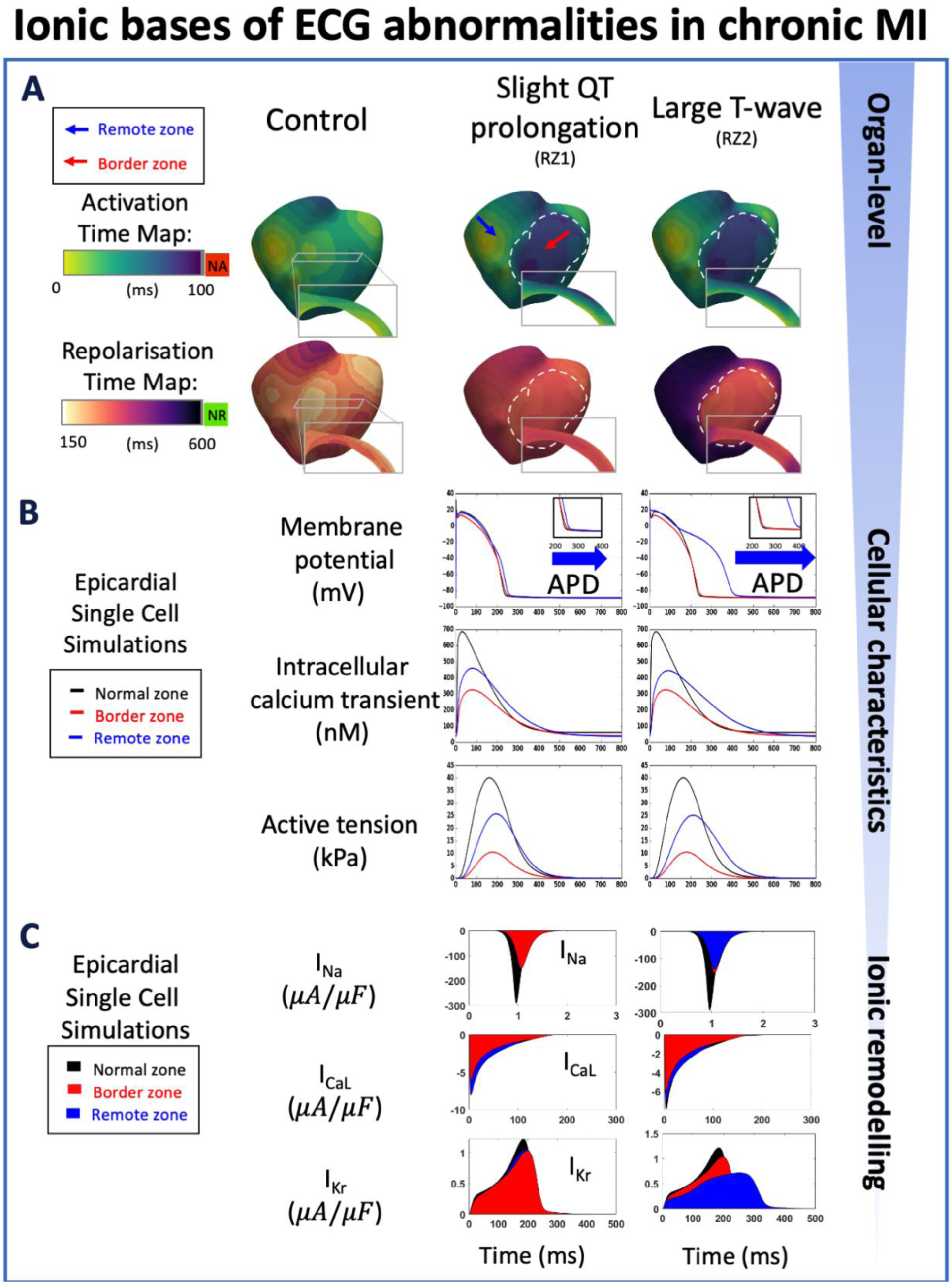
Multiscale explanation of QT and T-wave phenotypes in chronic MI. (A) Conduction delay in chronic border zone causes slight QT prolongation and large T-wave phenotypes, as well as large repolarisation dispersion in large T-wave. Red in activation map show regions of no activation (NA), green in repolarisation map show regions of no repolarisation (NR). (B) Varying degrees of action potential duration (ADP) prolongation in the remote zone (RZ) corresponding to extent of QT prolongation (blue arrows), with decreased calcium amplitude in remote and border zone of both phenotypes, and corresponding decrease in active tension generation. (C) As in acute MI, I_Na_, I_CaL_, and I_Kr_ remodelling underpin reduced conduction, reduced calcium amplitude, and degree of prolongation in action potential duration, respectively, in both chronic phenotypes.

Brugada phenocopy ECG phenotype was also observed in acute MI with BZ2 remodelling, causing regions of activation failure in the epicardial border zone (Figure 3A, the activation and repolarisation time maps), as well as delayed repolarisation in the BZ near the apex, caused by 29 ms of epicardial APD prolongation compared with control (Figure 3B, membrane potential). These were caused by strong inhibitions of sodium, calcium and potassium ionic currents in the BZ (Figure 3C, the second column of I_Na_, I_CaL_, I_Kr_, and SM Table S4 Acute BZ2). In this simulation, electrical activation in the infarcted region was preserved despite the conduction block in the BZ because of higher expressions of the L-type calcium channel in the mid-myocardium (see SM Figure S11).

Acute MI with upright T-waves corresponded to a comparable transmural repolarisation gradient as in control with BZ3 ionic remodelling (Figure 3A, the repolarisation time map). In this case, the lack of potassium current inhibition accounted for the lack of ECG manifestations (Figure 3C, the third column of I_Kr_, and SM Table S4 Acute BZ3).

### 3. In chronic post-MI, variable T-wave width can be explained by the extent of repolarisation dispersion between border zone and remote zone

In chronic MI, global QT prolongation was due to APD prolongation in the remote myocardium (Figure 4, repolarisation time maps and membrane potentials). Recovery of the upright T-wave in the anterior leads (compared to acute MI) was due to a recovery of the transmural repolarisation gradient (Figure 4A, repolarisation time maps, and SM Figure S10), given the milder I_Kr_ inhibition in the border zone (Figure 4B, the first column of I_Kr_, and SM Table S4 Chronic BZ). Furthermore, T-wave duration in this stage was mainly determined by the gradient between remote and border zone repolarisation times (Figure 4C, repolarisation time maps), where more severe APD prolongation in the RZ led to larger repolarisation gradients and, consequently, larger T-wave duration and amplitude (Figure 2B). Specifically, there was an APD difference of 157 ms between remote and border zone cell models for the large T-wave case versus only 12 ms for the slight QT-prolongation case, which accounts for the differences in T-wave peak-to-end duration (162 ms vs 72 ms) and QT intervals (565 ms vs 380 ms) between these two cases.

### 4. LVEF failed to indicate the extent of post-MI repolarisation dispersions

In our simulations, acute MI ionic remodelling yielded mildly reduced LVEF in the baseline ventricular models (43% to 47%) compared with control (53%) (Figure 5A). LVEF reductions were caused by contractile dysfunction (Figure 5A, active tension) due to lowered calcium amplitude in BZ (Figure 3B, intracellular calcium transient), which was directly caused by inhibitions of I_CaL_ in all acute phenotypes (Figure 3C, I_CaL_, and SM Table S4). A complete loss of contractile function in the BZ resulted in a more severe reduction in LVEF in acute post-MI (to 40% for all acute phenotypes in SM Figure S9). Stroke volumes of the left and right ventricles were well-matched in control conditions (1 mL difference, see SM Table S9), and introducing myocardial infarction caused a decrease of stroke volume in the left ventricles in both acute and chronic MI (see SM Table S9).

**Figure 5:**
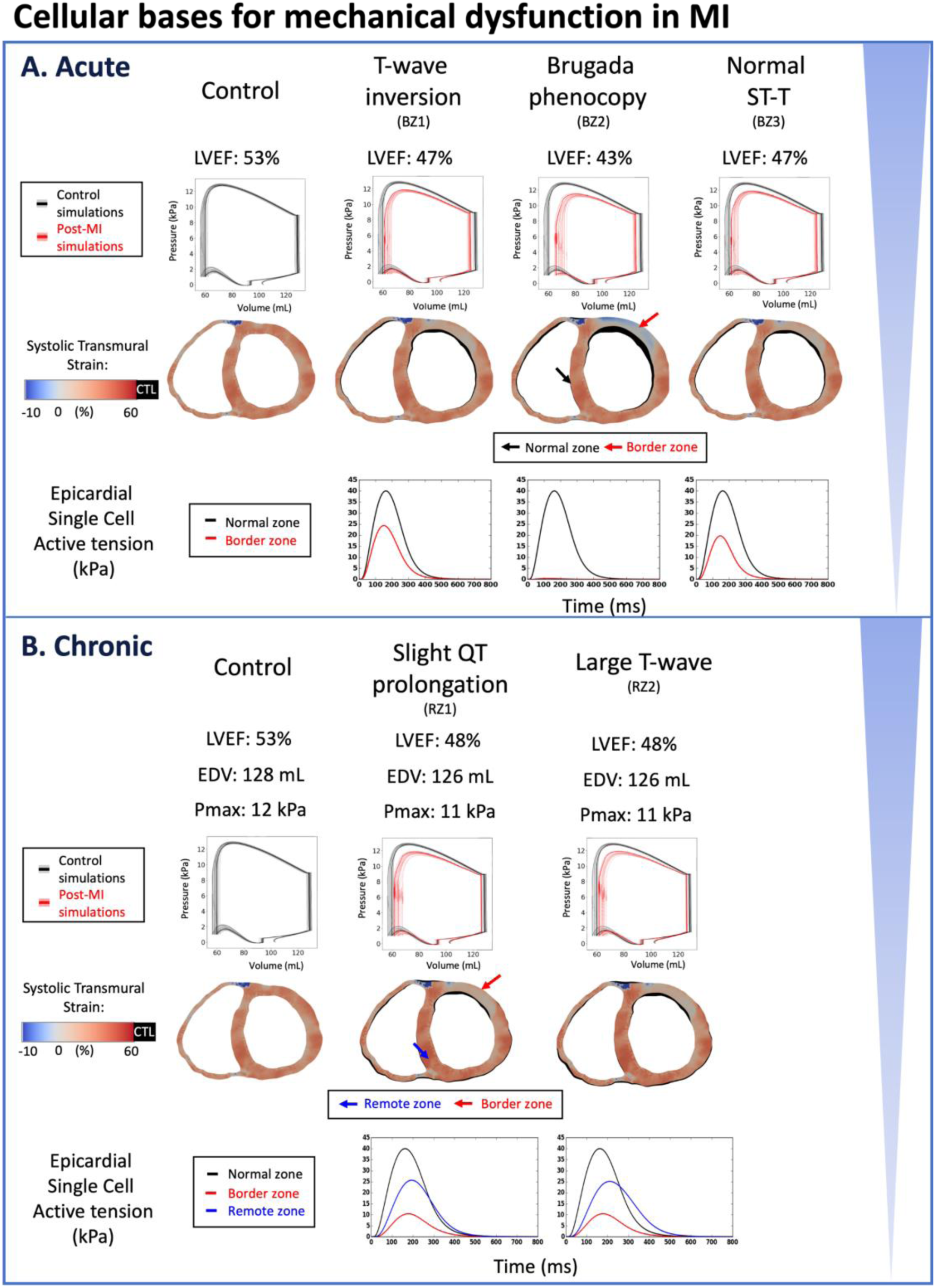
Reduced LVEF and heterogeneous systolic deformation caused by BZ ionic remodeling in both acute and chronic post-myocardial infarction. Pressure-volume loops are shown in black (control) or red (post-MI) traces for the baseline model, and in gray (control) or pink (post-MI) traces for the population of models. (A) Reduced LVEF in all acute phenotypes due to reduced active tension amplitude in the border zone (BZ1∼3). Brugada phenocopy shows the lowest LVEF due to activation block and loss of contractile function in the border zone in addition to reduced active tension amplitude in the activated border zone due to ionic remodelling. Reduced contractile function in infarct and border zone results in infarct thinning and bulging in systole. Systolic cross section of control simulation shown in black (CTL) with post-MI cross-sections superimposed. (B) Reduced LVEF in both chronic phenotypes due to reduced active tension amplitude in remote zone (RZ) and border zones, independent of the extent of QT prolongation (RZ1, RZ2). Scar stiffening helped to reduce infarct bulging. Systolic cross-section of control simulation shown in black (CTL) with post-MI cross-sections superimposed.

Activation failure in Brugada phenocopy caused loss of contractility in the affected regions in the BZ (Figure 3A, activation and repolarisation time maps) and resulted in a more severely reduced LVEF to 43% with a substantial non-contracting region with significant wall thinning (Figure 5A, systolic wall thinning). Acute MI with no observable ECG abnormalities can still have reduced mechanical function, as measured by an LVEF of 47% (Figure 5A, the last column). Comparison between this phenotype and the inverted T-wave phenotype shows that the mechanical dysfunction can be dissociated from repolarisation gradient and T wave abnormalities.

Chronic post-MI simulations showed mild reduction in peak systolic pressure (by 1 kPa) and some reduction in LVEF (Figure 5B), which were unaffected by differences in repolarisation gradients. This is because there is a consistent reduction in calcium transient amplitude in the remote zone that is independent of the extent of APD prolongation (Figure 4B, intracellular calcium transient and membrane voltage). Therefore, both the acute and chronic electromechanical simulation results showed that the post-MI repolarisation dispersion were not reflected by LVEF.

For both acute and chronic post-MI, simulations done using the population of ventricular models showed similar changes to the PV loop as the baseline model across variabilities in ionic current conductances.

### 5. T-wave alternans and abnormal wave propagation are caused at fast pacing by cellular alternans and EADs, without reduced LVEF at resting heart rate

Increased incidence of T-wave alternans is commonly observed in post-MI patients, and abnormally propagating waves generated from post-MI electrophysiological heterogeneity can trigger lethal arrhythmic events. T-wave alternans were reproduced in the ventricular chronic MI simulations at fast pacing (Figure 6A), with RZ2 remodelling, and their mechanisms were revealed through analysis of the high spatio-temporal resolution of simulation data. Figure 6A shows upright T-wave morphology and preserved LVEF of 49% at resting rate of 75 bpm (CL= 800 ms). However, at fast rates (120 bpm, CL = 500 ms), significant beat-to-beat ST and T-wave morphology alterations were observed. This was due to large alternans seen in mid-myocardial single cell simulations of the remote zone at CL of 500 ms (Figure 6C, green trace), with EAD-driven alternans. These results support the importance of stress tests, since alternans in APD and T-wave can occur at fast heart rates with no sign of LVEF abnormalities at resting heart rate. This is consistent with reports that T-wave alternans under supine bicycle exercise testing was found to be predictive of arrhythmic event after acute post-MI (Ikeda *et al*., 2000).

**Figure 6:**
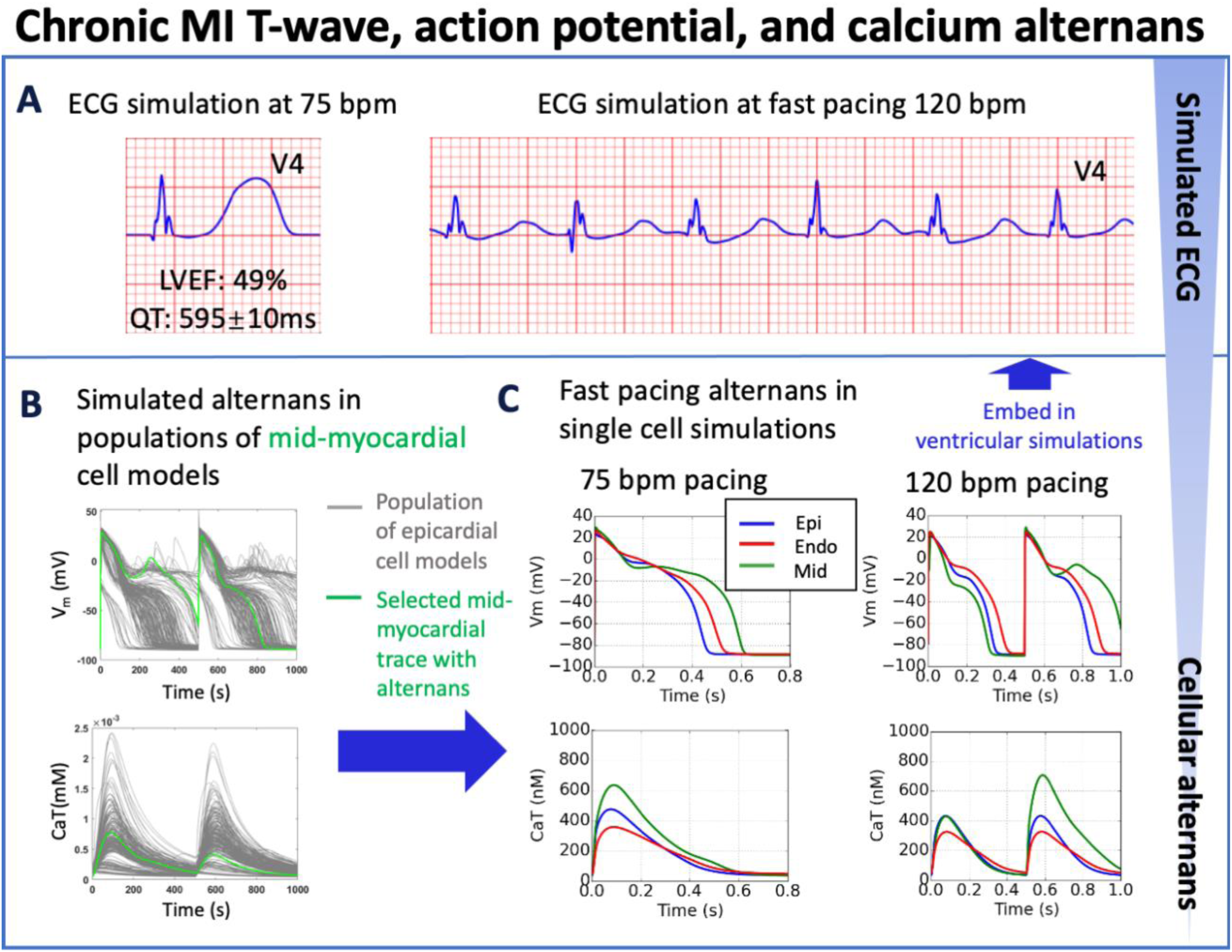
T-wave alternans in simulations underpinned by APD and calcium alternans at fast pacing (120 bpm), albeit with preserved left ventricular ejection fraction (LVEF=49%) at rest (75 bpm) for the chronic MI phenotype. (A). (B) Simulated APD and calcium traces in midmyocardial population of models with remote zone 2 (RZ2) remodelling. (C) Large action potential and calcium transient alternans were caused by EADs in simulations at 120 bpm with midmyocardial cells affected by RZ2 ionic remodelling (green traces, representative example at 75 versus 120 bpm). A single cell model (in green) was selected from the population of models (in grey) for embedding into the remote region for ventricular simulations.

Simulations with the population of virtual cardiomyocytes models revealed that in addition to the EAD-driven alternans (SM Figure S14), classical calcium-driven alternans were also observed in the population of cell models (SM Figure S12 and S13). The key ionic remodelling underlying calcium-driven alternans include enhanced CaMKII activity and slower calcium release, as well as suppressed SERCA pump activity in the chronic MI, which are consistent with previous studies (Livshitz & Rudy, 2007; Zhou *et al*., 2016; Tomek *et al*., 2018) (SM Figure S15). I_KCa_ enhancement in the chronic MI suppressed alternans generation (detailed analysis provided in SM7, Figures S16 to S17). The median of calcium amplitude was larger in the alternans models than in the non-alternating post-MI models (SM Figure S19), in agreement with the preserved LVEF in the simulations. We did not simulate the effect of this classical calcium-driven alternans on the ECG because the higher pacing rate at which this phenomenon occurs requires the model to include beta-adrenergic inotropic effects to preserve realistic systolic mechanical function.

Another case of chronic MI simulation showed prolonged QT interval of 669 ms and preserved LVEF of 49% at 75 bpm (Figure 7A). At 120bpm, in this case, ECG reflected chaotic activity (Figure 7A, fast pacing simulation) and loss of coordinated mechanical function. Abnormal electrotonic waves were caused by large repolarization dispersions.

**Figure 7:**
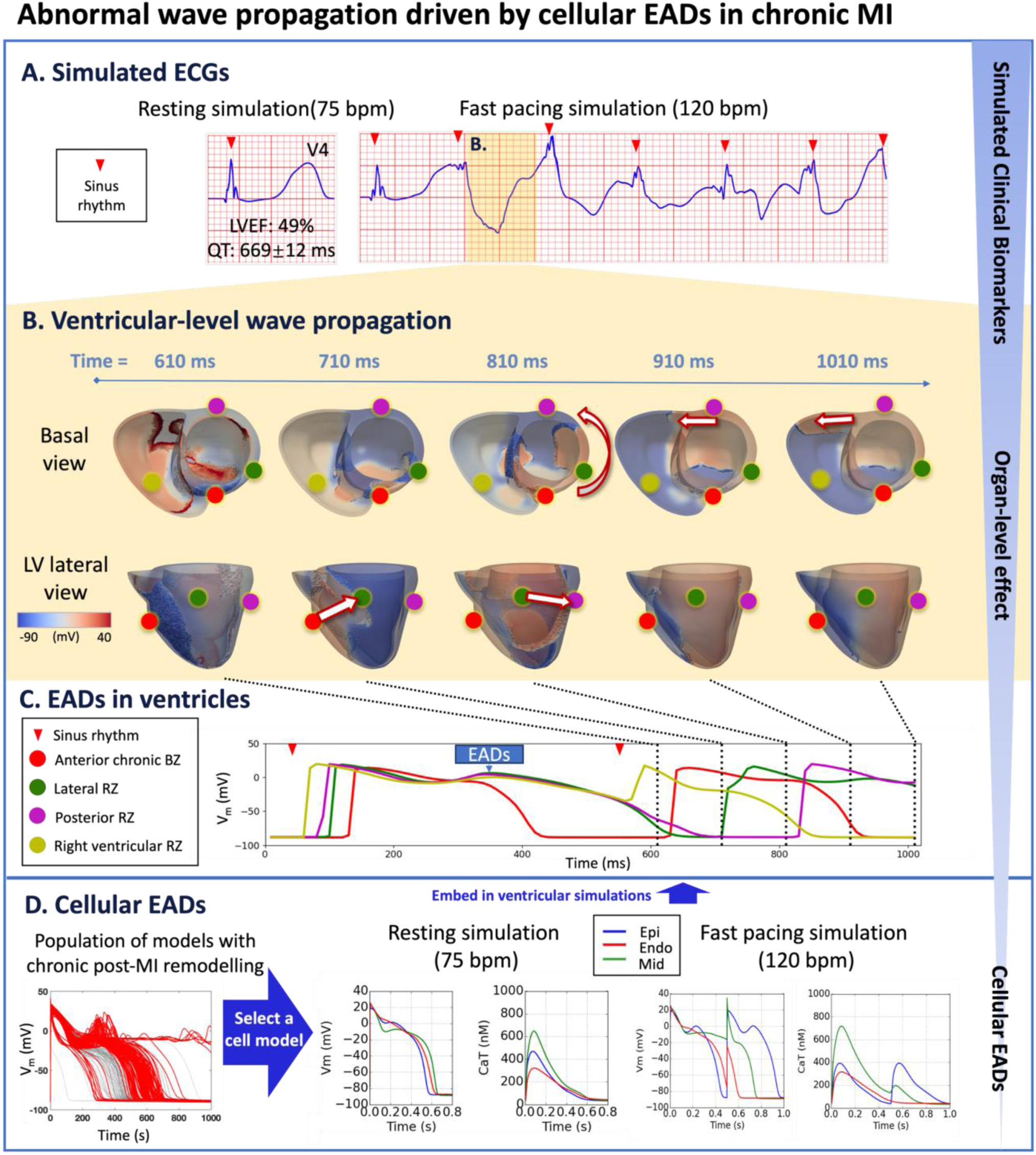
Prolonged QT and preserved LVEF at rest can manifest as severely abnormal ECG at fast heart rates in chronic MI with RZ2. (A), due to electrotonically-triggered EADs across the border zone (B). In (B), membrane potential changes for the first 1010 ms of fast pacing simulation, showing ectopic wave generation driven by electrotonic gradient at 710 ms (arrow from red dot to green dot in lateral view), and anticlockwise propagation of ectopic wave starting at 810 ms (anticlockwise arrow in basal view, and arrow from green dot to purple dot in lateral view). Ectopic wave propagates towards the right ventricle via the posterior side at 910 ms (arrow in basal view) and at 1010 ms (arrow in basal view). (C) Local action potential at anterior (red), lateral (green), posterior (purple), and right ventricular (yellow) sites. (D) A population of models demonstrating chronic remote zone 2 (RZ2) remodelling in promoting EADs. A representative example was selected from the population of models that showed EAD and was embedded in ventricular simulations.

When isolated cells that showed EADs were embedded in the RZ of a ventricular simulation at fast pacing, we saw ectopic wave propagation. This was because the EADs in the RZ generated conduction block, which enabled a large repolarisation gradient to form between the BZ and RZ, thereby leading to ectopy (Figure 7B). By the end of the first heartbeat (500 ms), the anterior epicardial BZ was fully repolarised (the red trace), but the RZ showed EADs (green, purple and yellow traces). This means that after the second beat stimulus, at 610 ms (Figure 7B), the long APD of the RZ prevented full activation in the postero-lateral LV (Figure 7B, green and purple marker) while the BZ APD was shorter than RZ (Figure 7B, red marker) and so could be fully activated. This led to a significant membrane potential gradient between the BZ (red marker) and RZ (green marker), leading to ectopic wave generation due to injury current caused by electronic effects at 710 ms in the boundary between BZ and RZ (Figure 7C, compare red and green traces)(Dutta *et al*., 2016). This ectopic wave propagated in an anti-clockwise fashion when viewed from the base from 810 to 910 ms (Figure 7B, basal view). In the right ventricle at 610 ms, incomplete repolarisation caused a smaller action potential to be elicited by the sinus stimulus (Figure 7C, yellow trace), thus allowing successful propagation of the ectopic wave from the left to the right ventricle from 910 ms to 1010 ms (Figure 7B).

This episode illustrated that EADs present in the epicardial remote zone cell model (Figure 7D) resulted in large APD dispersion between BZ and RZ, which functioned as the trigger of ectopic wave propagation due to electrotonic gradients. Therefore, the prolonged global QT interval with large T-wave duration and amplitude in leads facing the infarct can be indicative of the risk of large repolarization dispersion, while the LVEF can be preserved at rest.

Spontaneous EADs were frequently observed in the chronic MI cellular population of models (Figure 7D population of models, SM Figure S20), due to less I_CaL_ inhibition in the chronic MI (left, SM Table S4). The key underlying combination of ionic remodelling for EAD generation include the inhibition of I_Kr_ and the enhancement of I_NaL_, which facilitate reactivation of I_CaL_ (SM Figure S21). Additional contribution of baseline I_CaL_, I_Kr_, and I_NCX_ conductances are consistently observed in all chronic EAD populations (Table S11). Similarly, the cellular models that showed EADs also had larger calcium amplitude (SM Figure S22), suggesting a preserved LVEF.

## DISCUSSIONS

In this study, human electromechanical modelling and simulation enables quantification of the contribution of electrophysiological abnormalities to clinical phenotypes in post-MI patients, from ionic to whole-organ dynamics (summarised in Table 1). The credibility of the human electromechanical models and simulation results is supported by their consistency with experimental and clinical data from ionic dynamics to ECG and LVEF biomarkers in healthy, acute and chronic post-MI conditions. Diverse clinical ECG phenotypes are reproduced in the simulations with different degrees of experimentally reported ionic remodelling for acute and chronic MI; their signature on the LVEF is however weak, with only a small reduction observed. The simulated clinical ECG and LVEF phenotypes were found to be consistent across physiological variabilities in ionic conductances in the baseline electrophysiological model. Key findings include:

1. In acute MI, T-wave inversion, Brugada phenocopy, and QT prolongation were explained by reversed transmural dispersion of repolarisation in the border zone and infarct, activation failure in the border zone, and increased repolarisation time in the border zone, respectively.
2. In chronic MI, large T-wave duration and amplitude reflects large repolarisation dispersion between remote and border zones, and global QT prolongation was caused by AP prolongation in the remote zone.
3. Reduction in LVEF is ubiquitous across acute and chronic post-MI phenotypes and can be explained by decreased intracellular calcium amplitude and activation failure (in the case of Brugada phenocopy). This effect was independent of changes in the dispersion of repolarisation.
4. Interestingly, fast pacing simulations with T-wave alternans or abnormal propagation driven by cellular alternans or EADs both showed preserved LVEF at resting heart rate, highlighting the fact that preserved LVEF at rest does not guarantee low arrhythmic risk.

**Table 1:** Linking clinical ECG and left ventricular ejection fraction (LVEF) phenotypes to tissue heterogeneities and subcellular ionic remodelling in acute and chronic post-myocardial infarction.

| Clinical Phenotypes | Tissue or Cell Level phenomena | Corresponding Post Infarction Ionic Remodelling |
| --- | --- | --- |
| Acute MI T-wave inversion in ECG | Reversed transmural repolarisation gradient due to delayed repolarization in the epicardial border zone | Inhibition of potassium currents in the border zone |
| Acute MI<br>Brugada<br>phenocopy in<br>ECG | Delayed repolarisation, as well<br>as a small region of activation<br>failure in the epicardial border<br>zone | Strong inhibitions of sodium,<br>calcium and potassium ionic<br>currents in the border zone |
| Chronic MI<br>upright tall T-waves in ECG | Large repolarisation time<br>gradient between remote and<br>border zones caused by more<br>severe delay of repolarization in<br>the remote zone | More severe potassium channel<br>suppression in the remote zone |
| Chronic MI T-wave alternans | Cellular repolarization alternans<br>or early afterdepolarization | Suppressed SERCA and<br>augmented CaMKII activity for<br>alternans; Enhanced late sodium<br>current and suppressed hERG<br>current for early<br>afterdepolarization |
| Acute MI<br>reduction in<br>LVEF | Reduced calcium amplitude<br>and/or regional conduction block | Inhibitions of calcium and<br>sodium currents |
| Chronic MI<br>reduction in<br>LVEF | Reduced calcium amplitude | Decreased SERCA activity |

Collectively, our results show the proarrhythmic post-MI electrophysiological dispersions caused by cellular remodelling of ionic currents are reflected in QT and T wave morphology biomarkers rather than in LVEF, which questions the use of LVEF as the dominant biomarker in clinical risk stratifications.

### 1. Acute MI T-wave inversion and Brugada phenocopy are caused by reversed transmural repolarisation gradient and regional conduction abnormality

Three distinct types of T-wave morphology were generated by our acute MI biventricular simulations: T-wave inversion, Brugada-phenocopy and normal upright T-wave. We obtained them by applying three types of acute BZs in ventricular simulations, considering both APD prolongation and shortening, as a reflection of the variable experimental results (Mendonca Costa *et al*., 2018). Collectively, our results highlighted the importance of investigating the implications of the various degrees of experimentally-reported ionic remodelling to explain phenotypic variability of patients with MI. T-wave inversion is a commonly observed feature in the acute MI patients, and is commonly associated with arrhythmic risk (Tikkanen *et al*., 2015). Here we showed the reversed transmural repolarisation gradient caused by APD prolongation in the epicardial BZ accounted for this phenotype. The link between transmural repolarisation gradient and T-wave polarity has been reported previously (Okada *et al*., 2011) and is consistent with our results. Brugada-phenocopy was also observed in some acute MI patients (Anselm *et al*., 2014), and our simulation results showed it could be a reflection of regional conduction abnormality combined with APD prolongation. Although some animal experiments showed acute MI BZ APD shortening (Mendonca Costa *et al*., 2018), we showed the prolongation of BZ APD was underlying the QT prolongation, T-wave inversion and Brugada phenocopy in the leads facing the infarct (Supplementary Material Figure S7 showing lead dispersions), which is consistent with the QTc prolongation observed in the anterior leads of acute anterior infarction patients (Guaricci *et al*., 2018). Apart from the above, normal T-waves and QT intervals were also commonly observed in patients post percutaneous coronary intervention, which can be reproduced when the post-MI repolarisation dispersion was small between BZ and NZ (BZ3). It is worth noting that, in addition to having a mild border zone remodelling as shown in BZ3, a silent ECG signature can also be due to a reduced transmural extent of the infarct, as has been shown in previous computational studies (Loewe *et al*., 2018; Wang *et al*., 2021). Therefore, T-wave inversion, Brugada phenocopy and QT prolongation occur in the leads facing the infarct can be useful biomarkers indicating bigger repolarisation dispersions and/or larger transmural extent in the acute MI.

### 2. Wide and tall T-wave is explained by large repolarisation dispersions between BZ and RZ in healed post-MI hearts

Our simulated chronic ECGs recapitulated the recovery of T-wave polarity observed in patients after a period of healing. This was achieved through the recovery of the transmural repolarisation gradient caused by the milder I_Kr_ inhibition in the chronic BZ (Hegyi *et al*., 2018). Experimental studies in different species showed inconsistent results regarding the chronic BZ APD (Mendonca Costa *et al*., 2018). Our chronic BZ remodelling produced slightly longer APD than the NZ, which is consistent with observations in healed human BZ (Dangman *et al*., 1982). However, in minipigs, these remodelling caused shorter BZ APD than in NZ (Hegyi *et al*., 2018). This interesting discrepancy between minipigs and human may be due to the different balance of ionic currents across species, which showed the benefits of human electrophysiology models in overcoming the inter-species differences.

The two types of T-wave morphologies in the simulated chronic ECGs corresponded to different extents of RZ APD prolongations, which were commonly observed in healed RZ of post-MI animals (Hegyi *et al*., 2018), and in failing human myocytes (Li *et al*., 2004). The substantial RZ APD prolongation was reflected as global QT prolongation in all leads, and the large APD dispersion between chronic BZ and RZ generated wide and tall T-waves in the precordial leads facing the infarct (Supplementary Material Figure S8 for global and dispersed ECG characteristics). Previous simulation studies also found the T-wave amplitude and area were proportional to the dispersion of repolarisation (Arteyeva & Azarov, 2017). Therefore, these results demonstrated that in patients with global QT prolongation, leads with bigger T-wave amplitudes could reflect increased local heterogeneity in repolarisation.

### 3. T wave alternans and severely abnormal ECGs at fast pacing are caused by alternans and EADs in chronic infarction

Post-MI ionic remodelling promoted alternans generation, which resulted in T-wave alternans in simulated ECGs, consistent with the higher incidence of T-wave alternans reported in post-MI patients (Martin *et al*., 2009). Two types of repolarisation abnormalities were observed in our post-MI models: EADs and alternans. One crucial mechanism promoting alternans behaviour at the cellular level is the increased activity of CaMKII, observed in the acute MI BZ (Hund *et al*., 2008b), as well as in the hypertrophied and failing myocardium (Anderson *et al*., 2011). Enhanced CaMKII phosphorylation may preserve the contractility of the heart through the phosphorylation of phospholamban and the L-type calcium channels, but increased RyR phosphorylation by CaMKII resulted in prolonged RyR opening as well as the enhancement of spontaneous calcium sparks, which can contribute to alternans and triggered arrhythmias (Maier & Bers, 2007).

Generation of EADs due to chronic post-MI ionic remodelling, such as I_Kr_ inhibition and I_NaL_ enhancement, was consistent with previous studies (Coppini *et al*., 2013). We also found that both repolarisation reserve remodelling (I_NaL_ and I_Kr_) and calcium system remodelling (J_up_ and CaMKII) are important for the EAD-driven alternans (details provided in SM9). EADs in the RZ can create large repolarisation dispersion in the ventricle, facilitating abnormal electrotonic wave propagations. Similar re-entrant waves caused by electrotonic gradients were also observed in previous studies of acute ischemia (Ridley *et al*., 1992; Dutta *et al*., 2016; Boukens *et al*., 2021).

### 4. LVEF should be combined with QT and T-wave characteristics for arrhythmic risk stratification

In this study, we observe non-structurally-induced reductions of LVEF in the both the acute and the chronic post-MI stages. At both stages, ventricles with different extents of repolarisation dispersion may have similar LVEF because they have similar degrees of calcium reduction (acute MI T-wave inversion vs normal ST-T, and chronic MI two cases). Models with inducibility of T-wave alternans and arrhythmia at fast pacing rates may present with a preserved LVEF at resting heart rates (Figure 6-7). Our cellular level results also showed models with inducibility of repolarisation abnormalities, such as alternans and EADs, tended to have more preserved CaT magnitudes at rest rates. Therefore, these phenomena all support the fact that preserved LVEF measured at rest does not guarantee low arrhythmic risk.

A recent clinical study of post-MI patients with preserved LVEF showed defibrillators are needed in those patients with electrophysiological risk factors, such as prolonged QTc, increased T-wave alternans, to prevent sudden cardiac death (Gatzoulis *et al*., 2019). Consistently, we also found post-MI alternans and EADs can present as alternans of T-wave morphology and prolonged QT intervals. In addition to the global ECG changes, our simulation results also showed increased local repolarisation dispersion can be reflected in the leads facing the infarct: inverted T-wave and prolonged QT in the acute MI, and wide and tall T-wave in the chronic MI. Therefore, we suggest the consideration of these signs as markers of high arrhythmic risk.

### 5. Limitations

The main goal of this study is to investigate phenotypic variability in ECG and LVEF biomarkers arising from post-MI ionic remodelling. We have shown that the relationship between phenotypic variabilities and ionic remodelling remains consistent across physiological ranges of variation of the ionic conductances in the baseline cell model. Other sources of variability were not considered in this study including: heart anatomy, location and timing of the early activates sites, calcium sensitivity, conduction velocity. These could all modulate quantitatively the findings but we do not anticipate strong implications in the findings. The effect of variability in location and size of the scar on the ECG has been explored elsewhere(Li *et al*., 2024). LVEF reduction in clinical cases are more significant than in our simulations due to factors other than ionic remodelling: RZ structural remodelling and elevated myocardium stiffness, and abnormalities in anatomy. However, these effects have been well-documented elsewhere and this study serves to elucidate the non-structural mechanisms that underpin LVEF reduction that is linked to electrophysiological remodelling and arrhythmic risk. The basal plane in our simulation was fixed in space, which was necessary due to the segmented geometry from clinical MRI and to prevent unphysiological motion at the truncated basal plane. Despite this limitation, our conclusions regarding the relative comparisons of mechanical dysfunction are likely to still hold.

In addition, post-MI ionic remodelling can be modulated by other acute and chronic factors such as autonomic modulation (beta-adrenergic effects), inflammation, cell death, and metabolic remodelling, which can be explored in future work. The limitations in this study call for a need for personalised digital twins to be generated in the future to facilitate a better understanding of the interaction between structural remodelling and electrophysiological alterations.

## 6. CONCLUSIONS

Human-based electromechanical simulations reveal ionic mechanisms underlying T-wave inversion, Brugada phenocopy, and upright T-wave in acute post-MI, as well as the upright T-wave, QT prolongation and T-wave alternans in the healed chronic MI. In acute MI, while the potassium current reduction in the border zone was implicated for all phenotypes, the more severe ECG abnormalities in the Brugada phenocopy implicates additional remodelling for the sodium and calcium currents, which were also key factors in reduced mechanical function. In chronic MI, the degree of QT prolongation and the generation of pro-arrhythmic injury currents were directly related to the severity of potassium current remodelling in the remote region. In addition, late sodium current remodelling could be an important factor underpinning T-wave alternans in chronic MI through the promotion of EAD-driven alternans. Our results show that T-wave inversion, wide and tall T-wave, and QT prolongation in the leads facing the infarct are indicative of local dispersion of repolarisation, which is independent from the reduction of LVEF. Our simulation results suggest the utilisation of T-wave morphology, T-wave alternans and QT prolongation to improve risk stratification biomarkers even when the resting LVEF is preserved.

## Acknowledgements

The authors would like to acknowledge Dr Erica Dall’Armellina, Dr Arka Das, Dr Chris Kelly, and Dr Lei Wang, for discussions on the study.

## Preprint

This manuscript was first published as a preprint: Xin Zhou, Zhinuo Jenny Wang, Julia Camps, Jakub Tomek, Alfonso Santiago, Adria Quintanas, Mariano Vazquez, Marmar Vaseghi, Blanca Rodriguez (2022). [Clinical phenotypes in acute and chronic infarction explained through human ventricular electromechanical modelling and simulations]. bioRxiv. https://www.biorxiv.org/content/10.1101/2022.02.15.480392v3

