## Supplementary Material for "Clinical phenotypes in acute and chronic infarction explained through human ventricular electromechanical modelling and simulations"

### SUPPLEMENTARY MATERIALS

#### Table of Contents

SM1, SM2, SM3 – Multi-scale model descriptions: baseline cellular ventricular model (SM1), the baseline biventricular electromechanical model (SM2), and the ionic remodelling details for the border and remote zones in post-MI (SM3).

SM4, SM5 – Validation data (SM4) and sensitivity analysis (SM5).

SM6 – Supplementary results for simulations of ECG and pressure volume characteristics of post-MI phenotypes.

SM7, SM8 , SM9– Descriptions of the sub-cellular mechanisms of arrhythmia in the post-MI population of models.

#### SM1: Implementation of the calcium activated potassium current in the ToR-ORd model, and the generation of population of ToR-ORd-SK models.

As the calcium activated potassium current ($I_{KCa}$) were reported to be enhanced in heart failure, a new formulation of the $I_{KCa}$ was added into the ToR-ORd model based on published data(Chang *et al.*, 2013) to obtain an updated model named ToR-ORd-SK model. Due to the coupling of $I_{KCa}$ channels and the L-type calcium channels(Zhang *et al.*, 2018), the ratio of $I_{KCa}$ channels in the subspace was set to be the same as the L-type calcium channels in the model. The conductance of $I_{KCa}$ (gkca) was chosen to get a similar current density ratio between $I_{KCa}$ and $I_{Kr}$ as observed in minipig myocytes(Hegyi *et al.*, 2018*a*). The conductance of the background potassium current was scaled to 90% to adapt to the implementation of $I_{KCa}$. The formulation of this new $I_{KCa}$ current is the following:

gkca= 0.003;
ikcan=3.5;
kdikca=6.05e-04
Fraction$I_{KCa}$ss=0.8;
Fraction$I_{KCa}$i=1-Fraction$I_{KCa}$ss;

$$I_{KCa}\_ss = gkca \times \text{Fraction}I_{KCa}ss \times \frac{Ca_{ss}^{ikcan}}{Ca_{ss}^{ikcan} + kdikca^{ikcan}} \times (Vm - EK)$$

$$I_{KCa}\_i = gkca \times \text{Fraction}I_{KCa}i \times \frac{Ca_{i}^{ikcan}}{Ca_{i}^{ikcan} + kdikca^{ikcan}} \times (Vm - EK)$$

$I_{KCa}$=$I_{KCa}$_ss+$I_{KCa}$_i;

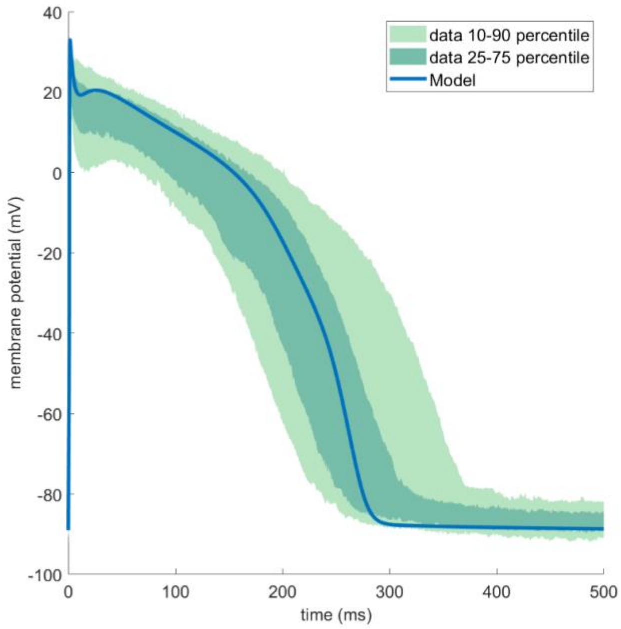


**Figure S1:** The ToR-ORd-SK model produced similar action potential (AP) traces as published human experimental data(O'Hara *et al.*, 2011).

An initial population of 500 human endocardial ventricular cell models was constructed based on the ToR-ORd-SK model by varying the conductances or magnitudes of $I_{Na}$, $I_{NaL}$, $I_{to}$, $I_{CaL}$, $I_{Kr}$, $I_{Ks}$, $I_{K1}$, $I_{NaCa}$, $I_{NaK}$, $J_{rel}$ and $J_{up}$ by up to ±50% using Latin Hypercube Sampling, and 253 models were accepted after pacing the models at 1 Hz and calibrated with human experimental data range in Table S1. The accepted endocardial parameter scaling factors were applied to the epicardial and midmyocardial baseline models to generate the corresponding epicardial/midmyocardial population of models, and eight models were discarded since they generate early afterdepolarizations (EADs) at 1 Hz in midmyocardial cells.

**Table S1:** Experimental ranges of AP and calcium transient (CaT) biomarkers used to calibrate the normal zone (NZ) endocardial population of models at a pacing cycle length (CL) of 1000ms based on human cardiomyocyte experiments(Coppini *et al.*, 2013; Britton *et al.*, 2017).

| Biomarkers at 1Hz | minimum | maximum |
|---|---|---|
| **Vmax (mV)** | 7 | 55 |
| **RMP (mV)** | -95 | -80 |
| **dvdtmax (mV/ms)** | 100 | 1000 |
| **APD90 (ms)** | 180 | 440 |
| **APD50 (ms)** | 110 | 350 |
| **APD40 (ms)** | 85 | 320 |
| **APD90-APD40 (ms)** | 50 | 150 |
| **CaTD90 (ms)** | 220 | 750 |
| **CaTD50 (ms)** | 120 | 420 |
| **CaTamp (mM)** | 2e-4 | 6e-4 |
| **CaTmax (mM)** | 2e-4 | 10e-4 |
| **CaTmin (mM)** | 0 | 4e-4 |

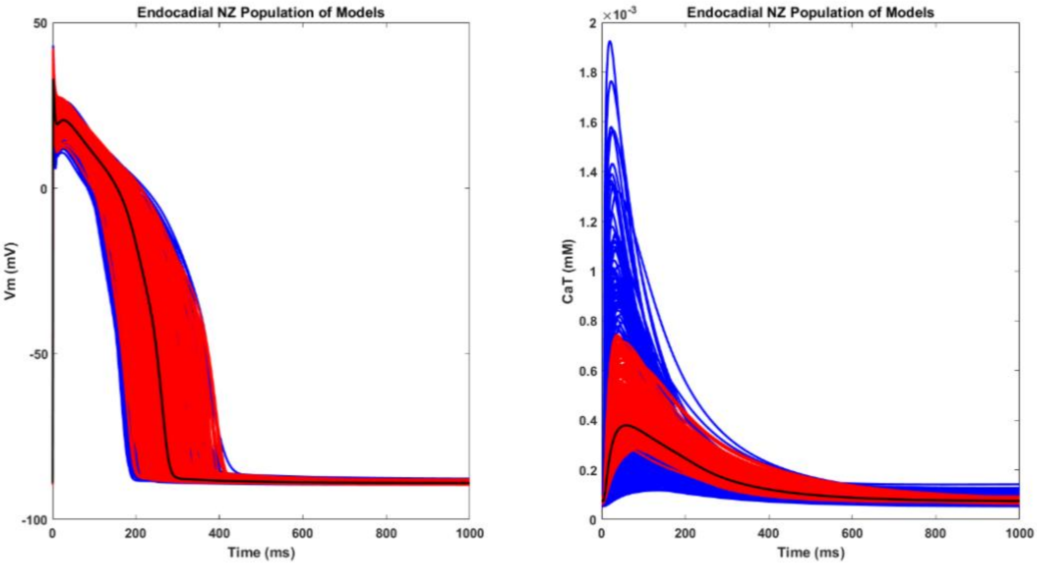

**Figure S2:** The AP and CaT traces of the population of NZ population of models. The blue and red traces are the initial and the accepted population, respectively. The black trace is the baseline endocardial model.

#### SM2: Human biventricular electromechanical models and calibrated simulation parameters

The parameters to reproduce the healthy baseline and infarcted models are presented in Tables S2 and Table S3.

The governing equations are listed here, and described in more detail elsewhere (Levrero-Florencio *et al.*, 2020):

$$\chi C_m \frac{\partial V}{\partial t} = \nabla \cdot (\boldsymbol{D}\nabla \mathrm{V_m}) + \chi(\mathrm{I_{ion}} + I_{stim})$$

$$\boldsymbol{D} = d_f \boldsymbol{f}\otimes\boldsymbol{f} + d_s \boldsymbol{s}\otimes\boldsymbol{s} + d_n \boldsymbol{n}\otimes\boldsymbol{n}$$

Where the orthotropic diffusivity tensor $\boldsymbol{D}$ describes different diffusivities along the fibre, sheet, and sheet normal $\boldsymbol{f}, \boldsymbol{s}, \boldsymbol{n}$ directions (values for $d_f, d_s, d_n$ given in Table S2), $\chi$ is the surface to volume ratio of the myocardial cell, $C_m$ the membrane capacitance, $V_m$ the transmembrane potential, and $I_{ion}, I_{stim}$ are the ionic channel currents and stimulus current, respectively.

The passive mechanical properties of the myocardium is described in the following strain energy density function $\psi$ in a nearly incompressible form of the Holzapfel Ogden orthotropic model (Holzapfel & Ogden, 2009):

$$\psi = \frac{K}{2}(J-1)^2 + \frac{a}{2b}\left(e^{b(I_1-3)} - 1\right) + \sum_{i=f,s}^{2} \frac{a_i}{2b_i}\left(e^{b_i(I_{4i}-1)^2} - 1\right) + \frac{a_{fs}}{2b_{fs}}\left(e^{b_{fs}I_{8fs}^2} - 1\right)$$

Where $K$ is the bulk modulus, and the strain invariants $I_1, I_{4f}, I_{4s}, I_{8fs}$ are invariants of the right Cauchy-Green strain tensor $\boldsymbol{C}$, as described below:

$$I_1 = tr\boldsymbol{C},\ \ I_{4f} = \boldsymbol{f}\cdot\boldsymbol{C}\boldsymbol{f}, I_{4s} = \boldsymbol{s}\cdot\boldsymbol{C}\boldsymbol{s}, I_{8fs} = \frac{1}{2}(\boldsymbol{f}\cdot\boldsymbol{C}\boldsymbol{s} + \boldsymbol{s}\cdot\boldsymbol{C}\boldsymbol{f})$$

And the values for ‘a’ and ‘b’ coefficients are given in Table S2.

The active stress $T_a$ produced through calcium-dependent cross-bridge cycling is calculated as:

$$T_a = T_{scale} \cdot h\frac{T_{ref}}{r_s}\left((\zeta_s + 1)S + \zeta_w W\right)$$

Where a scaling factor $T_{scale}$ is applied in the biventricular model to achieve a physiological left ventricular ejection fraction in control, and is given in Table S2, $h$ is a function of the fibre stretch ratio that describes the length-dependence of force production, $\zeta_s, \zeta_w$ are state variables describing distortion-decay, $W, S$ are the proportion of cross-bridges in the pre- or post-powerstroke states, respectively, and $r_s$ is a steady-state duty ratio of cross-bridge cycling. Details of this can be found elsewhere (Land *et al.*, 2017; Levrero-Florencio *et al.*, 2020).

**Table S2:** Calibrated electromechanical parameters for healthy baseline model, and modified parameters for post myocardial infarction models.

| **Name** | **Parameter** | **Value** | **Unit** |
|---|---|---|---|
| Healthy baseline electromechanical parameters | | | |
| diffusivity in fibre, sheet and sheet normal directions | $d_f$ | 0.00335 | cm/mS |
| | $d_s$ | 0.000723 | cm/mS |
| | $d_n$ | 0.000153 | cm/mS |
| active mechanics: scaling parameter for active tension | $T_{scale}$ | 12 | |
| bulk modulus | K | 12185000 | Ba |
| passive mechanics: exponential term in isotropic matrix, fibre, sheet and normal direction | a | 20000 | Ba |
| | b | 9.242 | |
| | $a_f$ | 30000 | Ba |
| | $b_f$ | 15.972 | |
| | $a_s$ | 20000 | Ba |
| | $b_s$ | 10.446 | |
| | $a_{fs}$ | 10000 | Ba |
| | $b_{fs}$ | 11.602 | |
| Scar and border zone diffusion parameters | | | |
| diffusivity in fibre, sheet and sheet normal directions | $d_f$ | 0.0012 | cm/mS |
| | $d_s$ | 0.00023 | cm/mS |
| | $d_n$ | 0.000003 | cm/mS |
| Scar mechanical parameters | | | |
| active mechanics: scaling parameter for active tension | $T_{scale}$ | 0 | |
| bulk modulus | K | 12185000 | Ba |
| passive mechanics: exponential term in isotropic matrix, fibre, sheet and normal direction | a | 200000 | Ba |
| | b | 9.242 | |
| | $a_f$ | 300000 | Ba |

| | | | |
|---|---|---|---|
| | $b_f$ | 15.972 | |
| | $a_s$ | 200000 | Ba |
| | $b_s$ | 10.446 | |
| | $a_{fs}$ | 100000 | Ba |
| | $b_{fs}$ | 11.602 | |

The pressure boundary condition on the left and right endocardial surfaces are controlled using the following set of piece-wise functions:

1) *Initialisation*. Both ventricles were firstly inflated to an initial pressure $P_0$ to reach a loaded resting endocardial volume, $V_0$.
2) *Active inflation.* The pressure in both ventricular chambers is linearly increased to the end diastolic pressure $P_{endd}$ over duration of $t_{diastole}$. This phase mimics the atrial contraction phase of diastolic filling and it is considered the first phase in the cardiac cycle because it follows directly from sinoatrial stimulus.
3) *Isovolumetric contraction.* Endocardial activation marks the beginning of this phase where stimulated myocytes begin generating active tension and the endocardial pressure (P) is allowed to increase such that the chamber volume (V) is kept approximately constant through the use of penalty terms:

$$dP = -\frac{1}{C_p}dV - \frac{1}{C_v} \cdot \frac{dV}{dt}$$

where $C_p$ and $C_v$ are the penalty terms for volume difference and volume rate, respectively. Furthermore, while $C_v$ is a user defined constant, $C_p$ is defined as:

$$C_p = \frac{P}{V}$$

4) Ejection. This phase is triggered when the ventricular pressure exceeds the arterial pressure, ${P_{art}}_0$. A two-element Windkessel model is used to model the blood pressure of both the systemic and pulmonary circulation systems during ejection:

$$C\frac{dP_{art}}{dt} + \frac{P_{art}}{R} = -\frac{dV}{dt}$$

where C and R are the compliance and impedance of the circulation systems.
5) *Isovolumetric relaxation.* This phase is triggered by the reversal of ventricular volume change, i.e. $\frac{dV}{dt} > 0$. Symmetrically with phase 2), here the pressure is allowed to decrease while the volume is kept constant.
6) *Passive filling.* This phase is triggered when ventricular pressure is lower than a threshold pressure $P_{post}$. During this phase the myocyte active tension is allowed to return to resting state and the volume is allowed to return to the initialised value $V_0$ through:

$$dP = -\frac{1}{C_p}(V - V_0) - \frac{1}{C_v} \cdot \frac{dV}{dt}$$

where ventricular volumes were calculated at each time step using the divergence theorem (Levrero-Florencio *et al.*, 2020). Here, both the $C_p$ and $C_v$ parameters are user defined and have been selected to allow full recovery of the initialised volume before the end of the cycle length.

**Table S3:** Parameters for boundary conditions and phase control at resting heart rate and fast pacing (in brackets).

| Name | Parameter | LV | RV | Unit |
|---|---|---|---|---|
| Pericardial stiffness | $K_{epi}$ | 10000 | | Ba cm$^{-1}$ |
| Time to initial pressure | $t_0$ | 0.02 | 0.02 | s |
| Initial pressure | $P_0$ | 5000 | 5000 | Ba |
| Duration of passive diastolic filling | $t_{diastole}$ | 0.08 (0.03) | 0.08 (0.03) | s |
| Pressure at end of diastole | $P_{endd}$ | 15000 | 15000 | Ba |
| Arterial compliance | C | 0.00055908 | 0.00055908 | cm$^3$ Ba$^{-1}$ |
| Arterial resistance | R | 250 | 100 | Ba s cm$^{-3}$ |
| Aortic pressure | $P_{art0}$ | 90000 | 20000 | Ba |
| Pressure at end of isovolumetric relaxation | $P_{post}$ | 10000 | 10000 | Ba |
| Penalty parameters for isovolumetric contraction | $C_v$ | 1 | 1 | cm$^3$ s$^{-1}$ Ba$^{-1}$ |
| Penalty parameters for isovolumetric relaxation | $C_v$ | 0.2 | 0.2 | cm$^3$ s$^{-1}$ Ba$^{-1}$ |
| Penalty parameters for passive filling | $C_p$, $C_v$ | 0.1,0.3 | 0.1,0.3 | cm$^3$ Ba$^{-1}$, cm$^3$ s$^{-1}$ Ba$^{-1}$ |

**ECG simulation and biomarker calculation**

Simulated ECGs were evaluated at prescribed electrode locations on the torso surface using the pseudo-ECG method (Mincholé *et al.*, 2019), assuming the torso is an infinite volume conductor. The signals are normalised in the precordial leads according to the maximum amplitude, and they are then normalised separately for the limb leads. This addresses the limitation of this method in not being able to faithfully representing absolute amplitudes (Ogiermann *et al.*, 2021). Only the precordial leads were evaluated because the biomarkers calculated for these leads were the most reliable.

ECG biomarkers as listed in Table S7 were evaluated using a Python code as follows:

1) Separate each beat using the pacing R-to-R interval.
2) Resample the signal to 1000 Hz.
3) For each beat, use the voltage at end of each beat to offset the signal.
4) Evaluate the first and second order derivative of the voltage signal, apply filter on the

derivatives using python package scipy.signal.lfilter with filtering parameters tuned manually to achieve best results for steps 4), 5) and 6). Normalize the absolute value of the filtered signal using the maximum absolute value.

5) Identify beginning of QRS by searching from the beginning of signal and finding the first time at which the first derivative becomes higher than the threshold 0.01*max(V)/30.
6) Identify the end of QRS manually.
7) Identify the end of T wave by searching from the end of signal and finding the first time point at which the first derivative becomes higher than the threshold 0.01*max(V)/30.
8) Isolate the signal segment from the end of QRS to the end of T wave and evaluate the peak absolute value to find T-wave peak and identify its timing.
9) Isolate the signal segment from the end of QRS to the peak of T wave. Evaluate mean dV using a window width of 3, then, beginning at the end of the QRS, the onset of T wave is identified as the time step at which the second derivative of voltage first becomes larger than 0.12*max(V)/30.
10) Evaluate the required biomarkers using the landmark points identified.

An example of the output of this delineation method is as below, showing QRS duration, QT duration, and T-wave duration (units ms) are shown after the lead name. Due to low reliability of this method in delineating the QRS complex, only the QT duration and T wave characteristics are reported in the main manuscript.

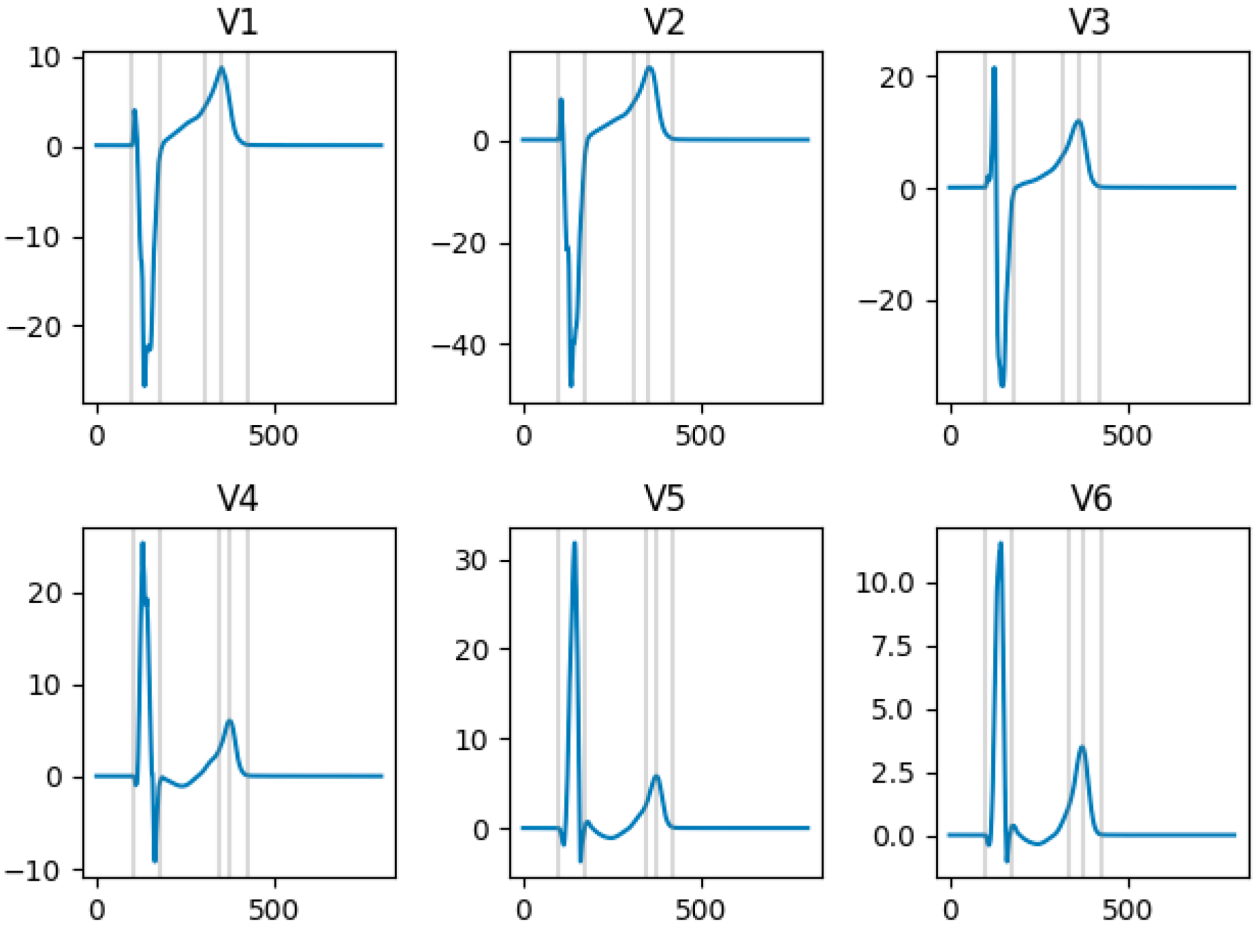
V1
V2
V3
V4
V5
V6
0
500

#### SM3: Border zone (BZ), remote zone (RZ) and scar ionic remodeling and their effects on AP and CaT biomarkers

For acute post-MI (within a week post-occlusion), three types of BZ remodelling (Acute BZ1-3) were considered based on previous modelling work and experimental canine data collected within 5 days post-MI(Hund *et al.*, 2008; Decker & Rudy, 2010; Arevalo *et al.*, 2016; Tomek *et al.*, 2017). For chronic post-MI, ionic remodelling measured from minipigs 5 months post-MI with heart failure were used to generate Chronic BZ (affecting only the BZ) and Chronic RZ1 (also affecting the remote myocardium). Another type of remodelling, Chronic RZ2, was established based on multiple experimental data from failing human cardiomyocytes(Schwinger *et al.*, 1999; Jiang *et al.*, 2002; Li *et al.*, 2004; Zicha *et al.*, 2004; Valdivia *et al.*, 2005; Maltsev *et al.*, 2007; Holzem Katherine M *et al.*, 2011; Chang *et al.*, 2013; Elshrif *et al.*, 2014; Gomez *et al.*, 2014; Hegyi *et al.*, 2018*b*; Høydal *et al.*, 2018). Furthermore, reduction of sodium current(Valdivia *et al.*, 2005) and SERCA(Jiang *et al.*, 2002), with enhanced CaMKII activity and slower calcium release(Hoch *et al.*, 1999; Maier & Bers, 2007) were also implemented in Chronic BZ, RZ1 and RZ2, as observed in human failing cardiomyocytes. The remodelling in the infarcted scar region was modelled as previously published(Wang *et al.*, 2021), which generated prolonged action potential than the normal zone as observed in the activation recovery interval (ARI) data of post infarction human and pigs(Vaseghi *et al.*, 2017; Srinivasan *et al.*, 2019).

**Table S4:** Ionic remodelling for the acute and chronic stage BZ and RZs.

| **Scaling** | **Acute BZ1** | **Acute BZ2** | **Acute BZ3** | **Chronic BZ** | **Chronic RZ1** | **Chronic RZ2** | **Infarct** |
|---|---|---|---|---|---|---|---|
| $G_{Na}$ | 0.4 (Hund *et al.*, 2008; Decker & Rudy, 2010) | 0.38 (Arevalo *et al.*, 2016) | 0.4 (Tomek *et al.*, 2017) | 0.43 (Valdivia *et al.*, 2005) | 0.43 (Valdivia *et al.*, 2005) | 0.43 (Valdivia *et al.*, 2005) | 0.4 |
| $G_{NaL}$ | | | | 1.275 (Hegyi *et al.*, 2018*b*) | 1.413 (Hegyi *et al.*, 2018*b*) | 2 (Valdivia *et al.*, 2005; Maltsev *et al.*, 2007) | |
| $G_{to}$ | 0.1 (Hund *et al.*, 2008; Decker & | | 0 (Tomek *et al.*, 2017) | | | 0.6 (Beuckelmann *et al.*, 1993; Li *et al.*, 2004) | 0 |

| | | | | | | | |
|---|---|---|---|---|---|---|---|
| | Rudy, 2010) | | | | | | |
| $G_{CaL}$ | 0.64 (Hund *et al.*, 2008; Decker & Rudy, 2010) | 0.31 (Arevalo *et al.*, 2016) | 0.64 (Tomek *et al.*, 2017) | 0.7 (Hegyi *et al.*, 2018*b*) | | | 0.64 |
| $G_{Kr}$ | 0.7 (Hund *et al.*, 2008; Decker & Rudy, 2010) | 0.3 (Arevalo *et al.*, 2016) | | 0.89 (Hegyi *et al.*, 2018*b*) | 0.87 (Hegyi *et al.*, 2018*b*) | 0.6 (Ambrosi *et al.*, 2013) | 0.7 |
| $G_{Ks}$ | 0.2 (Hund *et al.*, 2008; Decker & Rudy, 2010) | 0.2 (Arevalo *et al.*, 2016) | | | | 0.4 (Li *et al.*, 2004) | |
| $G_{K1}$ | 0.3 (Hund *et al.*, 2008; Decker & Rudy, 2010) | | 0.6 (Tomek *et al.*, 2017) | 0.76 (Hegyi *et al.*, 2018*b*) | | 0.6 (Beuckelmann *et al.*, 1993; Li *et al.*, 2004) | 0.6 |
| $G_{NaK}$ | | | | | | 0.6 (Schwinger *et al.*, 1999) | |
| $P_{Jup}$ | | | | 0.4 (Jiang *et al.*, 2002; Høydal *et al.*, 2018) | 0.4 (Jiang *et al.*, 2002; Høydal *et al.*, 2018) | 0.3 (Jiang *et al.*, 2002; Høydal *et al.*, 2018) | |

| | | | | | | | |
|---|---|---|---|---|---|---|---|
| $G_{KCa}$ | | | | 2 (Hegyi *et al.*, 2018*b*) | 2 (Hegyi *et al.*, 2018*b*) | 3.75 (Chang *et al.*, 2013) | |
| $G_{ClCa}$ | | | | 1.25 (Hegyi *et al.*, 2018*b*) | 1.25 (Hegyi *et al.*, 2018*b*) | 1.25 (Hegyi *et al.*, 2018*b*) | |
| aCaMK | | | 1.5 (Tomek *et al.*, 2017) | 1.5 (Hoch *et al.*, 1999; Hund *et al.*, 2008) | 1.5 (Hoch *et al.*, 1999; Hund *et al.*, 2008) | 1.5 (Hoch *et al.*, 1999; Hund *et al.*, 2008) | 1.5 |
| $Tau_{relp}$ | | | 6 (Maier *et al.*, 2003) | 6 (Maier *et al.*, 2003) | 6 (Maier *et al.*, 2003) | 6 (Maier *et al.*, 2003) | 6 |
| $G_{Cab}$ | | | 1.33 (Tomek *et al.*, 2017) | | | | 1.33 |

**Table S5:** Comparison of the simulated AP, CaT and active tension (Ta) biomarkers with post myocardial infarction (MI) acute stage canine experimental data and chronic stage human experimental data.

| | **Biomarkers** | **Experimental values** | **Simulated values** |
|---|---|---|---|
| Acute Post-MI Stage | Canine epi NZ APD (ms) (mean±SD） | 295±34 (Lue W M & Boyden P A, 1992)<br>210±15 (Gardner *et al.*, 1985)<br>219±39 (Spear *et al.*, 1983)<br>Overall: [180, 329] | 227 |
| | Canine epi BZ APD (ms) (mean±SD） | 346±60 (Lue W M & Boyden P A, 1992)<br>170±15 (Gardner *et al.*, 1985)<br>220±26 (Spear *et al.*, 1983)<br>Overall: [194, 406] | BZ1: 284,<br>BZ2: 256,<br>BZ3: 208 |
| | Canine epi BZ Systolic Cai EBZ/NZ (%) | 74% (Licata *et al.*, 1997) | NZ: 686,<br>BZ1: 517 (75%),<br>BZ2: 133 (20%),<br>BZ3: 457 (67%), |
| | Canine epi BZ Voltage Clamp Cai at 0mV EBZ/NZ | 53% (Pu *et al.*, 2000) | |
| | Canine Cell shortening EBZ/NZ % | 12% (Licata *et al.*, 1997) | NZ systolic Ta: 40,<br>BZ1: 24 (60%),<br>BZ2: 0.37 (1%),<br>BZ3: 20 (50%) |
| Chronic Post-MI Stage | Human Mid Systolic Cai Failing/Non-Failing (%) | 49% (Piacentino *et al.*, 2003) | |

| | | | |
|---|---|---|---|
| | Human Mid Systolic Cai MI/normal (%) 1 Hz | 37.5% (Høydal *et al.*, 2018) | NZ (800 ms CL):1219,<br>RZ1: 744 (60%),<br>RZ2: 608 (50%) |
| | Human Mid Diastolic Cai Failing/Non-Failing (%) | 96% (Piacentino *et al.*, 2003) | NZ (800 ms CL): 80,<br>RZ1: 37 (46%),<br>RZ2: 39 (49%)<br><br>NZ (500 ms CL): 86,<br>RZ1: 52 (61%),<br>RZ2: 62 (72%) |
| | Human Mid Diastolic Cai MI/normal (%) 1 Hz | 115% (Høydal *et al.*, 2018) | |
| | Human Mid Cell shortening MI/normal (%) 1 Hz | 33% (Høydal *et al.*, 2018) | NZ systolic Ta (800 ms CL): 65,<br>RZ1: 58 (89%),<br>RZ2: 45 (70%) |

**Table S6:** Simulated AP, CaT and Ta biomarkers from baseline NZ, BZ and RZ epi-, mid-, and endocardial single cell models. For the acute stage, the Acute BZ1 and BZ2 induced significant APD prolongation, while the BZ3 led to mild APD shortening. The Acute BZ1 and BZ3 had similar degree of reduction in systolic Ca and Ta, whereas the Acute BZ2 had more severe loss of contractility. For the chronic stage, the Chronic BZ1 and RZ2 had similar decrease in systolic Ca and Ta than control, but the RZ2 had more severe APD prolongation than the Chronic BZ.

| **Type** | **APD90 (ms)** | **CaTD90 (ms)** | **Diastolic Ca (nM)** | **Systolic Ca (nM)** | **Diastolic Ta (kPa)** | **Systolic Ta (kPa)** |
|---|---|---|---|---|---|---|
| Control (800 ms) | epi: 227, mid: 336, endo: 263 | epi: 298, mid: 333, endo: 336 | epi: 62.32, mid: 79.74, endo: 70.70 | epi: 686.27, mid: 1218.83, endo: 477.71 | epi: 0.06, mid: 0.10, endo: 0.07 | epi: 40.00, mid: 65.47, endo: 23.87 |
| Acute BZ1 | epi: 284, mid: 391, endo: 315 | epi: 289, mid: 339, endo: 325 | epi: 59.63, mid: 63.52, endo: 65.62 | epi: 517.13, mid: 691.30, endo: 342.98 | epi: 0.05, mid: 0.06, endo: 0.06 | epi: 24.33, mid: 44.78, endo: 10.46 |
| Acute BZ2 | epi: 256, mid: 373, endo: 341 | epi: 266, mid: 342, endo: 334 | epi: 45.11, mid: 57.08, endo: 57.98 | epi: 133.09, mid: 326.57, endo: 184.93 | epi: 0.03, mid: 0.05, endo: 0.05 | epi: 0.37, mid: 8.97, endo: 1.48 |
| Acute BZ3 | epi: 208, mid: 316, endo: 247 | epi: 256, mid: 295, endo: 288 | epi: 49.05, mid: 60.49, endo: 59.79 | epi: 457.49, mid: 834.58, endo: 352.71 | epi: 0.03, mid: 0.05, endo: 0.05 | epi: 19.66, mid: 52.08, endo: 11.18 |
| Chronic BZ | epi: 235, mid: 362, endo: 293 | epi: 419, mid: 444, endo: 474 | epi: 41.90, mid: 40.80, endo: 50.59 | epi: 324.87, mid: 499.97, endo: 285.67 | epi: 0.03, mid: 0.03, endo: 0.04 | epi: 10.44, mid: 31.40, endo: 7.76 |
| Chronic RZ1 | epi: 247, mid: 411, endo: 313 | epi: 426, mid: 462, endo: 478 | epi: 39.64, mid: 37.47, endo: 49.34 | epi: 459.72, mid: 744.07, endo: 387.91 | epi: 0.03, mid: 0.05, endo: 0.04 | epi: 25.67, mid: 57.56, endo: 18.40 |

| | | | | | | |
|---|---|---|---|---|---|---|
| Chronic RZ2 | epi: 392, mid: 591, endo: 467 | epi: 498, mid: 569, endo: 557 | epi: 39.77, mid: 38.61, endo: 49.51 | epi: 444.67, mid: 607.59, endo: 361.11 | epi: 0.03, mid: 0.09, endo: 0.05 | epi: 25.14, mid: 44.80, endo: 16.17 |
| Acute and (chronic) scar | epi: 250, mid: 366, endo: 295 | epi: 261, mid: 304, endo: 296 | epi: 49.17, mid: 62.49, endo: 59.70 | epi: 502.85, mid: 883.02, endo: 375.70 | epi: 0.03 (0), mid: 0.06 (0), endo: 0.05 (0) | epi: 24.23 (0), mid: 53.21 (0), endo: 13.28 (0) |
| Control (500 ms) | epi: 210, mid: 306, endo: 240 | epi: 265, mid: 276, endo: 298 | epi: 58.61, mid: 85.60, endo: 67.91 | epi: 760.33, mid: 1740.38, endo: 531.49 | epi: 0.38, mid: 1.81, endo: 0.33 | epi: 42.77, mid: 68.04, endo: 27.54 |
| RZ1 (500 ms) | epi: 226, mid: 296, endo: 271 | epi: 370, mid: 384, endo: 396 | epi: 47.10, mid: 52.52, endo: 65.07 | epi: 470.50, mid: 712.75, endo: 385.73 | epi: 0.50, mid: 2.04, endo: 0.51 | epi: 25.84, mid: 51.69, endo: 17.28 |
| RZ2 (500 ms) | epi: 316, mid: 395, endo: 366 | epi: 389, mid: 401, endo: 410 | epi: 50.88, mid: 61.53, endo: 65.99 | epi: 425.66, mid: 605.85, endo: 337.64 | epi: 0.52, mid: 1.94, endo: 0.46 | epi: 21.20, mid: 41.79, endo: 12.23 |

#### SM4: Validation using clinical and experimental data

The clinical ECGs in Figure 2 of main manuscript were extracted from the following:

Acute BZ1 comparison (T-wave inversion): PTB Diagnostic ECG Database (Goldberger *et al.*, 2000; Bousseljot *et al.*, 2009), patient number 014 who had anterior infarction. The ECG was taken 10 days after infarction.

Acute BZ2 comparison (Brugada phenocopy): Extracted from Figure 1B of a clinical paper describing Brugada phenocopy (Anselm *et al.*, 2014) and enhanced using bespoke python script. The patient had acute inferior ST segment elevation myocardial infarction with right ventricular involvement.

Acute BZ3 comparison (Normal ST-T): PTB Diagnostic ECG Database (Goldberger *et al.*, 2000; Bousseljot *et al.*, 2009), patient number 051 who had antero-septal infarction. The ECG was taken 10 days after infarction.

Chronic RZ1 comparison (Slight QT prolongation): PTB Diagnostic ECG Database (Goldberger *et al.*, 2000; Bousseljot *et al.*, 2009), patient number 042 who had antero-septal infarction. The ECG was taken at ~20 months follow up.

Chronic RZ2 comparison (Large T-wave): PTB Diagnostic ECG Database (Goldberger *et al.*, 2000; Bousseljot *et al.*, 2009), patient number 033 who had antero-septal infarction. The ECG was taken 3 months after infarction.

**Table S7:** Comparison of the ECG and mechanical biomarkers from biventricular electromechanical simulations against literature values at resting heart rate. QTc was calculated using Bazett's formula from the simulated QT intervals. Post-MI RVEF values were from ST-segment elevation myocardial infarction patients whose culprit and chronic total occlusion were not in the right coronary artery. VA: ventricular arrhythmia; VT: ventricular tachycardia; SDB: sleep disordered breathing. Our simulated ECG and mechanical biomarker values are mostly consistent with the clinically reported biomarker ranges.

| Biomarkers | Control | | Acute Stage Post-MI | | Chronic Stage Post-MI | |
|---|---|---|---|---|---|---|
| Electrophysiological Biomarkers | Literature | Simulation | Literature | Simulation | Literature | Simulation |
| QRS duration (ms) | 96 ± 9 in men, 85 ± 6 in women | 79±2 | 88±35(Yerra *et al.*, 2006) | 91±5, 95±9, 92±6 | Max 127±16 without VT Min 81 ± 15 without VT | 94±6, 93±5 |

| | | | | | | |
|---|---|---|---|---|---|---|
| | (Carlsson *et al.*, 2006) | | | | (Perkiömäki *et al.*, 1995) Max 137± 25 with VT Min 89± 20 with VT (Perkiömäki *et al.*, 1995) | |
| QTc interval (Bazett formula) (ms) | 350–440 (Johnson & Ackerman, 2009) | 360±1 | 423±50 without VA (Ahnve, 1985) 460±40 with VA (Ahnve, 1985) | 398±27, 415±4, 376±5 | Max 448±39 without VT Min 383±20 without VT (Perkiömäki *et al.*, 1995) Max 493±51 with VT Min 388±30 with VT (Perkiömäki *et al.*, 1995) | 430±4, 578±3 |
| Mechanical Biomarkers | Literature | Simulation | Literature | Simulation | Literature | Simulation |
| LVEDV (mL) | 142 ± 21 (SSFP-CMR) (Maceira *et al.*, 2006*a*) | 129 | 116±15(Uslu *et al.*, 2013) | 124-125 | 106±12 (Uslu *et al.*, 2013) | 126 |
| RVEDV (mL) | 144 ± 23 (SSFP-CMR) | 131 | 129±28 with SDB (Buchner *et al.*, 2015) | 131 | 143±29 with SDB | 133 |

| | | | | | | |
|---|---|---|---|---|---|---|
| | (Maceira *et al.*, 2006*b*) | | 132±28 without SDB (Buchner *et al.*, 2015) | | (Buchner *et al.*, 2015) 132±31 without SDB (Buchner *et al.*, 2015) | |
| LVESV (mL) | 47 ± 10 (SSFP-CMR) (Maceira *et al.*, 2006*a*) | 60 | 61±12 (Uslu *et al.*, 2013) | 65~72 | 52±10 (Uslu *et al.*, 2013) | 65 |
| RVESV (mL) | 50 ± 14 (SSFP-CMR) (Maceira *et al.*, 2006*b*) | 63 | 56±21 with SDB (Buchner *et al.*, 2015) 53±16 without SDB (Buchner *et al.*, 2015) | 64 | 58±21 with SDB (Buchner *et al.*, 2015) 51±15 without SDB (Buchner *et al.*, 2015) | 64 |
| LVEF (%) | 67 ± 4.6 (SSFP-CMR) (Maceira *et al.*, 2006*a*), 62 ± 7 (RNV) (Nemerovski *et al.*, 1982) | 53 | 48± 8 (Uslu *et al.*, 2013) | 43~47 | 52± 7 (Uslu *et al.*, 2013) | 48 |
| RVEF (%) | 48 ± 5 (RNV) (Nemerovski *et al.*, 1982) | 52 | 53.0 ± 7.1 (van Veelen *et al.*, 2022) | 51 | 55.9 ± 5.4 (van Veelen *et al.*, 2022) | 52 |

#### SM5: Sensitivity analysis of ECG and LVEF to electrophysiological heterogeneities, calcium sensitivity, and sheet active tension.

We performed the following sets of electromechanical simulations varying the apex-to-base and transmural heterogeneities, calcium sensitivity, and sheet active tension, in order to evaluate the effect of these on ECG morphology and LVEF.

The baseline apex-to-base gradient is evaluated by scaling the conductance of the slow delayed rectifier potassium current (GKs) according to:

$$sf = 0.2^{z_{scale}}$$

$$z_{scale} = 2z - 1$$

where z is the normalised longitudinal coordinate that is equal to zero at the apex and one at the base. This gives a scaling factor of 5 at the apex and 0.2 at the base (i.e., [0.2, 5]). We vary the magnitude of this gradient by changing the basis of exponential for the sf equation through the values [0.1, 0.2, 0.3], to achieve ranges of [0.1, 10], [0.2, 5], [0.3, 3.3] for the scaling factor (Figure S3A). In addition, we also reverse the gradient direction thus:

$$z_{scale} = -2z + 1$$

The effect of changes in range of the apex-to-base gradient is negligible in LVEF (Figure S3B) and ECG morphology (Figure S3C) except in leads V4 and V5, where the amplitude of the T-wave decreased with decreasing range of GKs scaling. A complete reversal of the gradient caused a more dramatic decrease in T-wave amplitude and reduction in T-wave duration apparent in V3 to V5 (Figure S3C, red line), with no changes in LVEF.

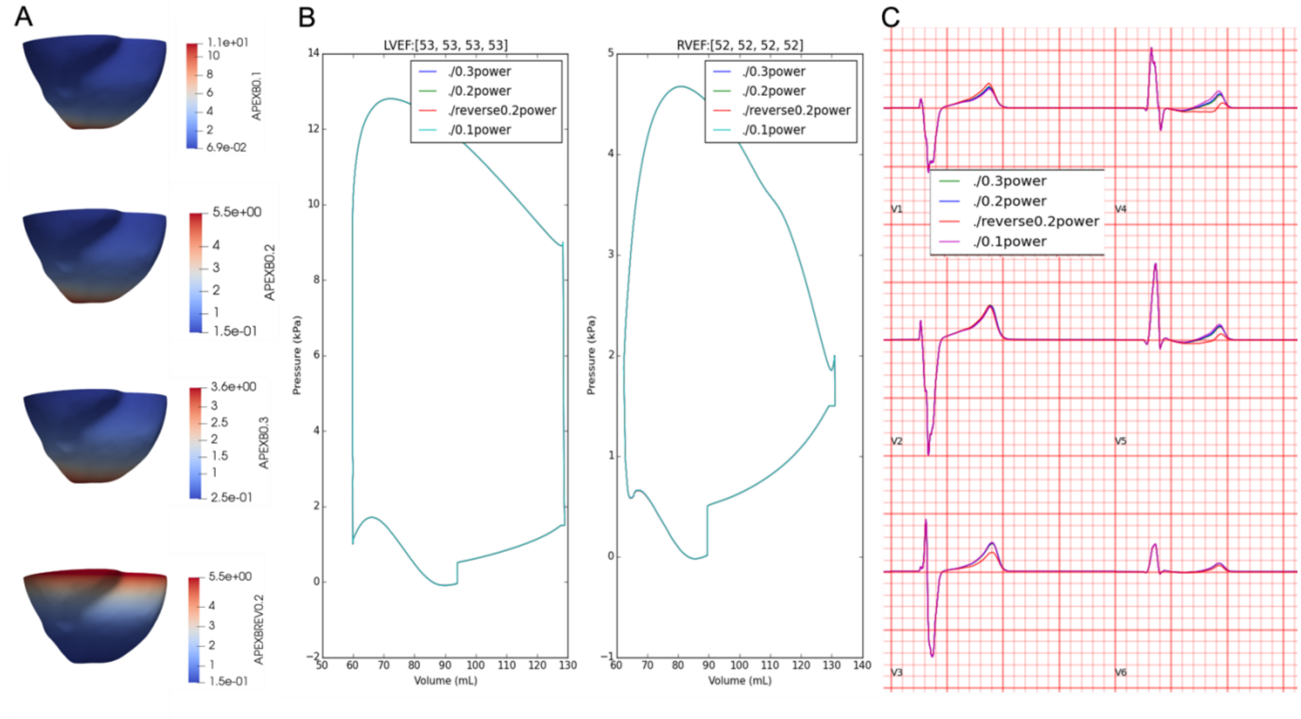


**Figure S3:** Effects of apex-to-base gradient (A) on pressure-volume, LVEF (B), and ECG morphology (C).

The baseline transmural heterogeneity has a 30%, 40%, and 30% split of endocardial, mid-myocardial, and epicardial cell types across the wall (Figure S4A). To modify this, we created two other transmural compositions where the mid-myocardial layer is removed, and the endo- vs. epicardial split is varied from 30%-70% to 50%-50% (Figure S4A). The removal of the mid-myocardial layer caused an increase in end systolic volume, a reduction in LVEF, and a decreased T-wave amplitude and increased QT interval across all precordial leads (compare green and blue). Increasing the proportion of epicardial cell type (compare red and blue) caused a reduction in T-wave amplitude and an increase in QT interval. The polarity of the T-wave remained unchanged.

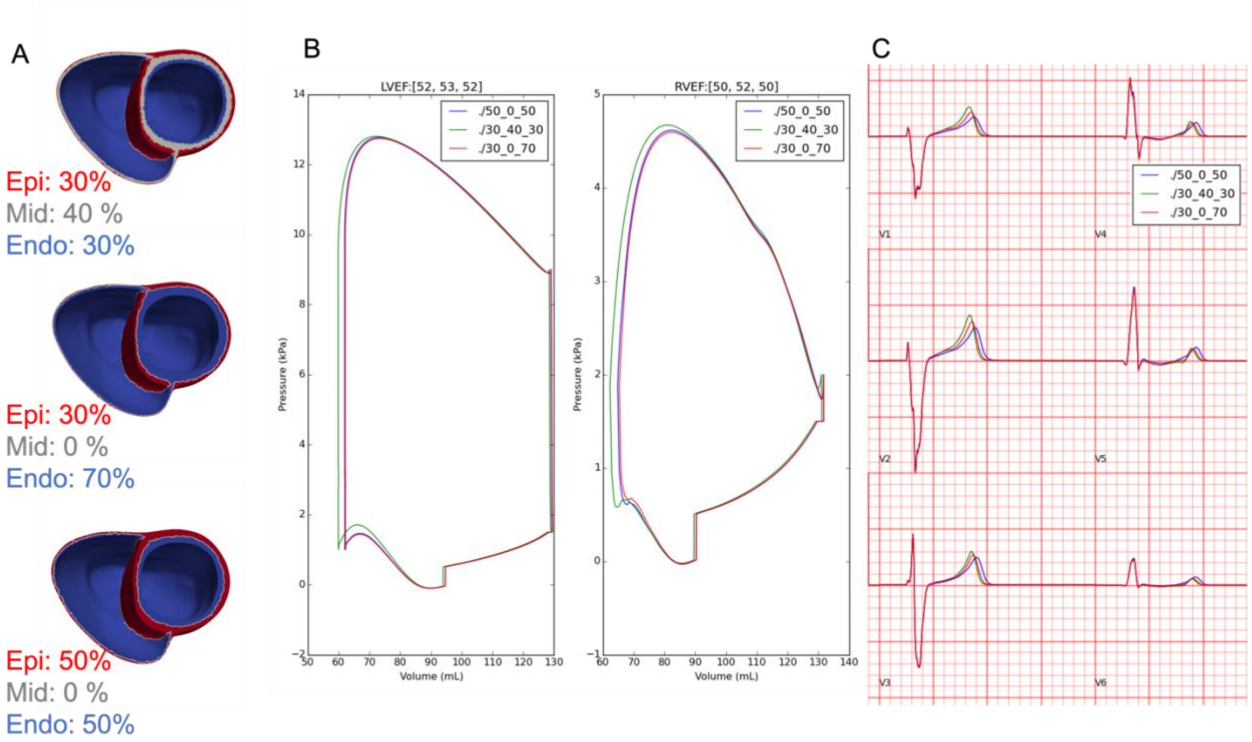


**Figure S4:** Effects of transmural electrophysiological heterogeneity (A) on pressure-volume, LVEF (B), and ECG morphology (C).

Additionally, we altered the calcium sensitivity of troponin binding in the excitation-contraction coupling (Ca50 parameter from the baseline value(Land *et al.*, 2017) of 0.805 $\mu M$ to a range of values: [0.5, 0.7, 1.0]. This had no effect on the ECG (Figure S5A). With increasing Ca50 values, there was an increase in end diastolic volume and end systolic volume, resulting in negligible changes to the LVEF, except in the case of 0.5, where the LVEF is reduced by 2% (Figure S5B).

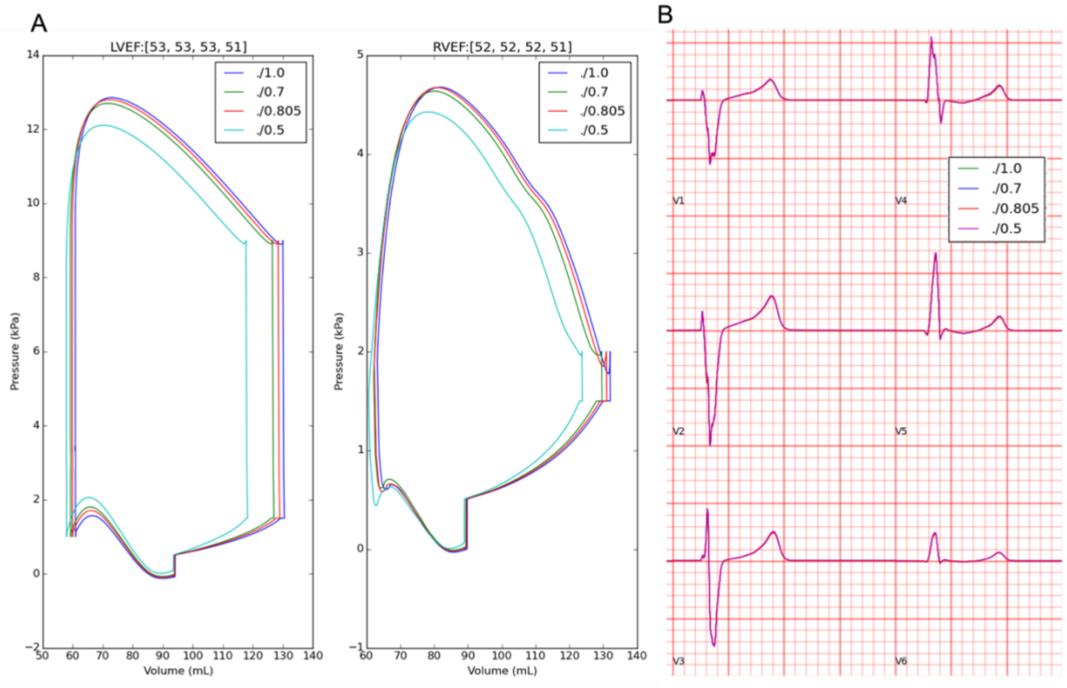


**Figure S5:** Effects of troponin calcium sensitivity on pressure-volume, LVEF (A), and ECG morphology (B).

Finally, we also altered the sheet-direction active tension from the baseline, where it is 30% of the fibre active tension, to a range of values: [0%, 50%, 60%]. With increasing percentage sheet activation LVEF increased (Figure S6A) with negligible effect on ECG morphology (Figure S6B). Sheet activation above and including 50% caused numerical instabilities that caused the simulation to terminate prematurely during the isovolumic contraction phase of the cardiac cycle.

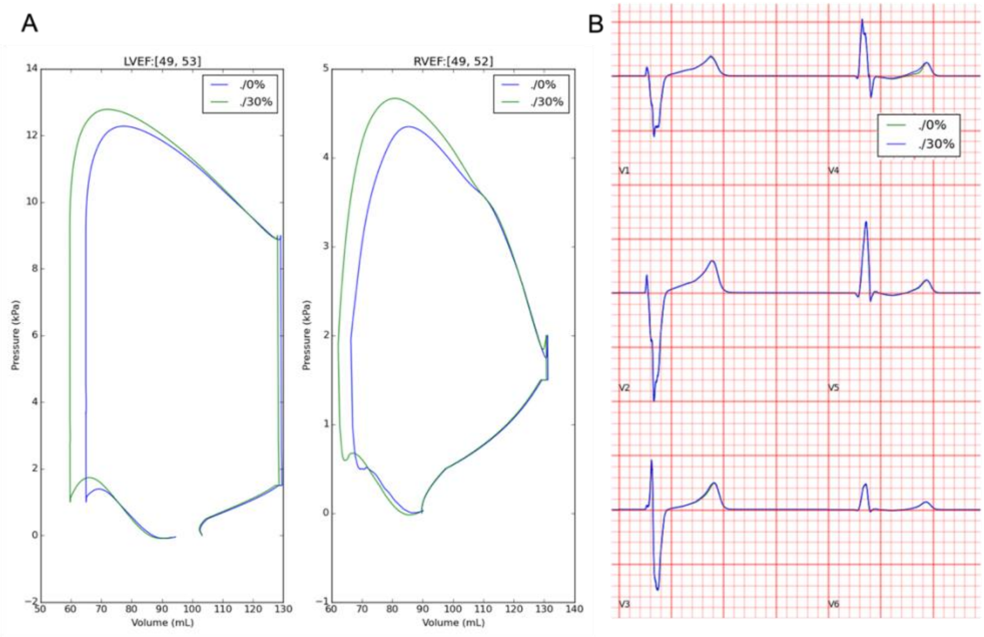


**Figure S6:** Effects of sheet activation percentage on pressure-volume, LVEF (A), and ECG morphology (B).

#### SM6: Biventricular electromechanical simulations of acute and chronic post-MI, ECG and pressure volume characteristics

Additional results of simulations for the acute and chronic post-MI phenotypes:

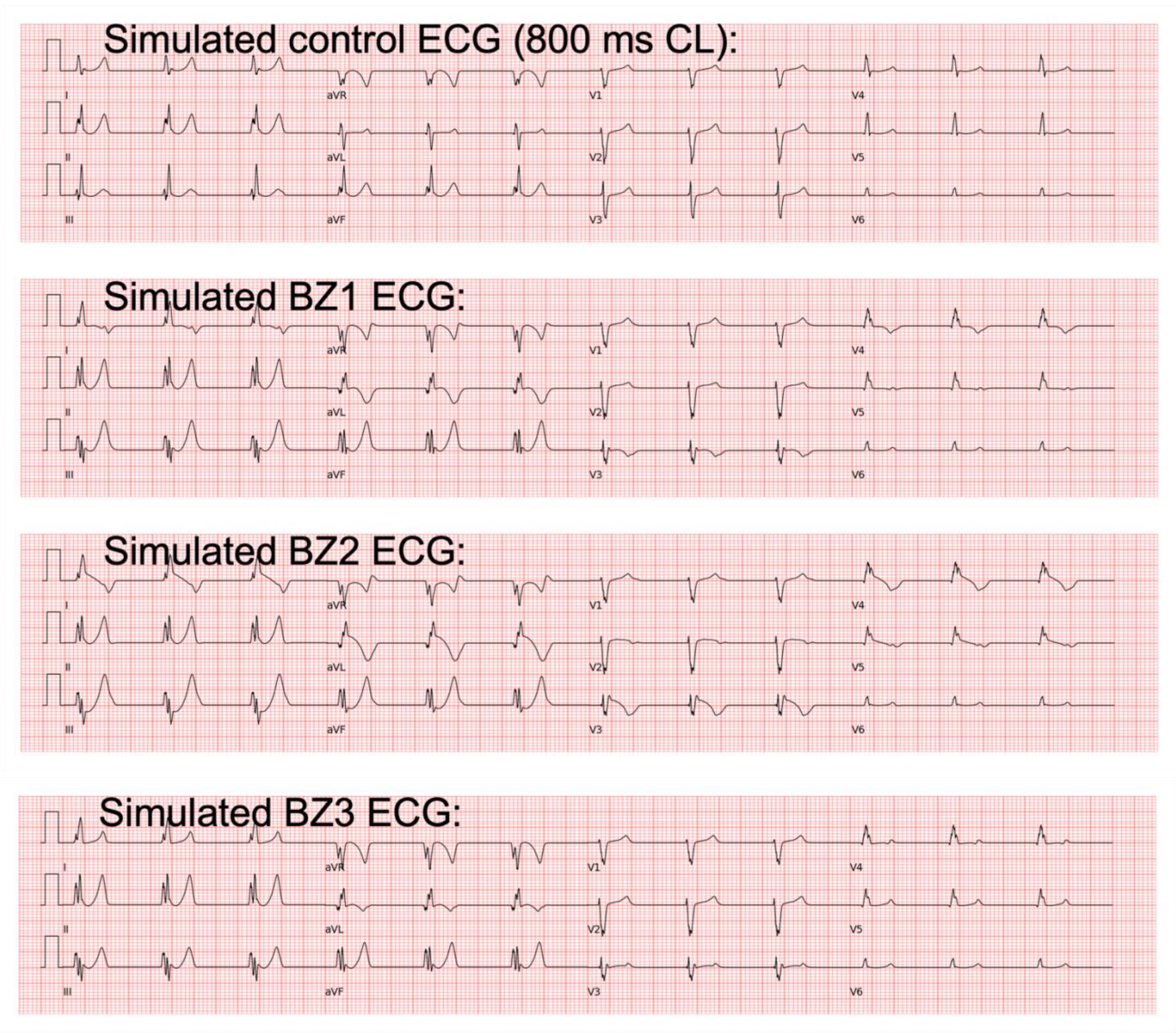


**Figure S7:** Simulated acute stage 12-lead ECGs for Acute BZ1-3. Acute BZ1 caused T wave inversion in precordial leads of V3 and V4, where the QT prolongation was more significant. Acute BZ2 caused Brugada phenocopy in leads V3-V5, while Acute BZ3 produced similar ECG morphology as the control case.

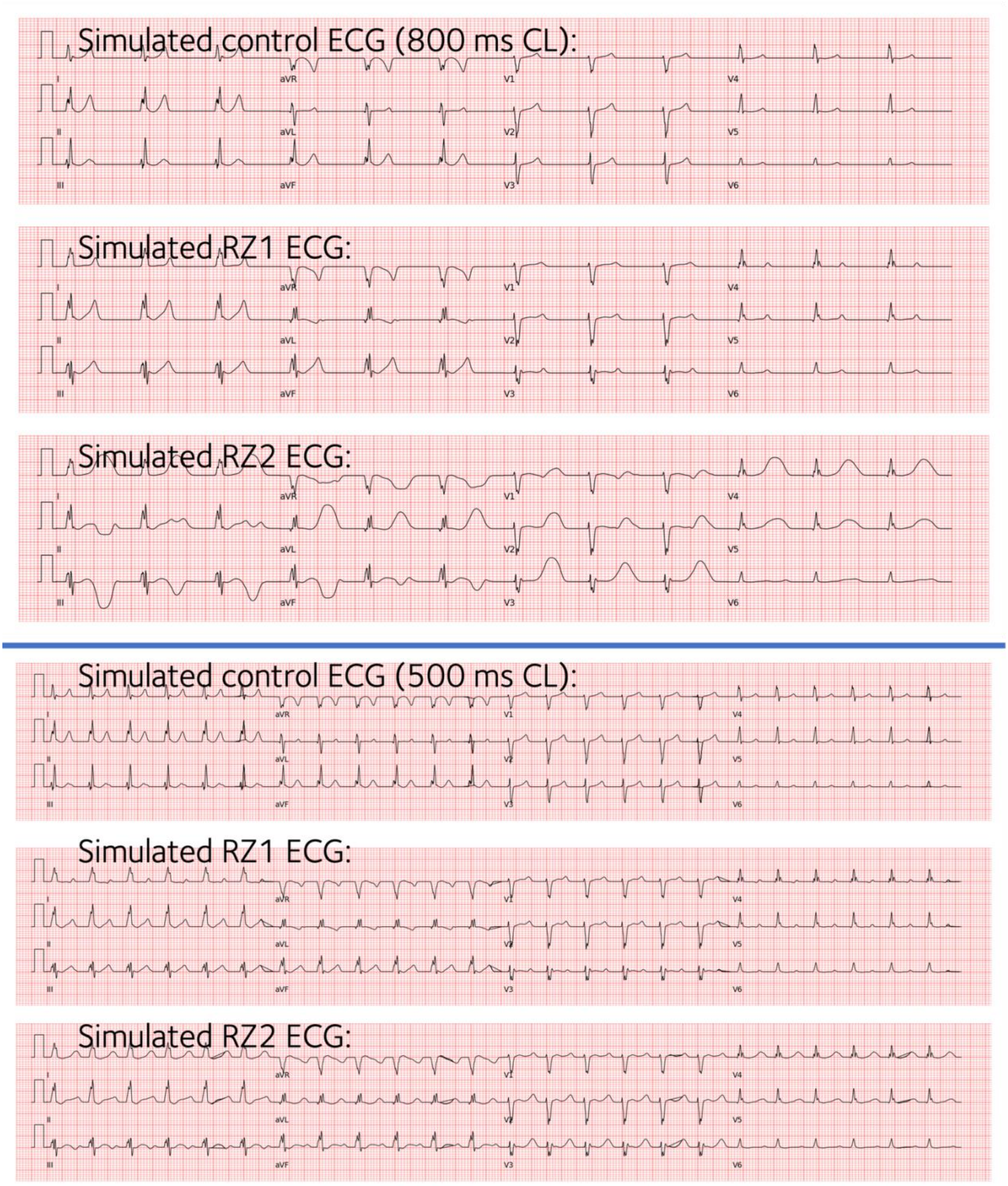


**Figure S8:** Simulated chronic stage 12-lead ECGs for Chronic RZ1 and Chronic RZ2, both combined with Chronic BZ. Both produced normal ECG morphology, and T waves are wider and taller in the anterior leads (V2-V4) of Chronic RZ2.

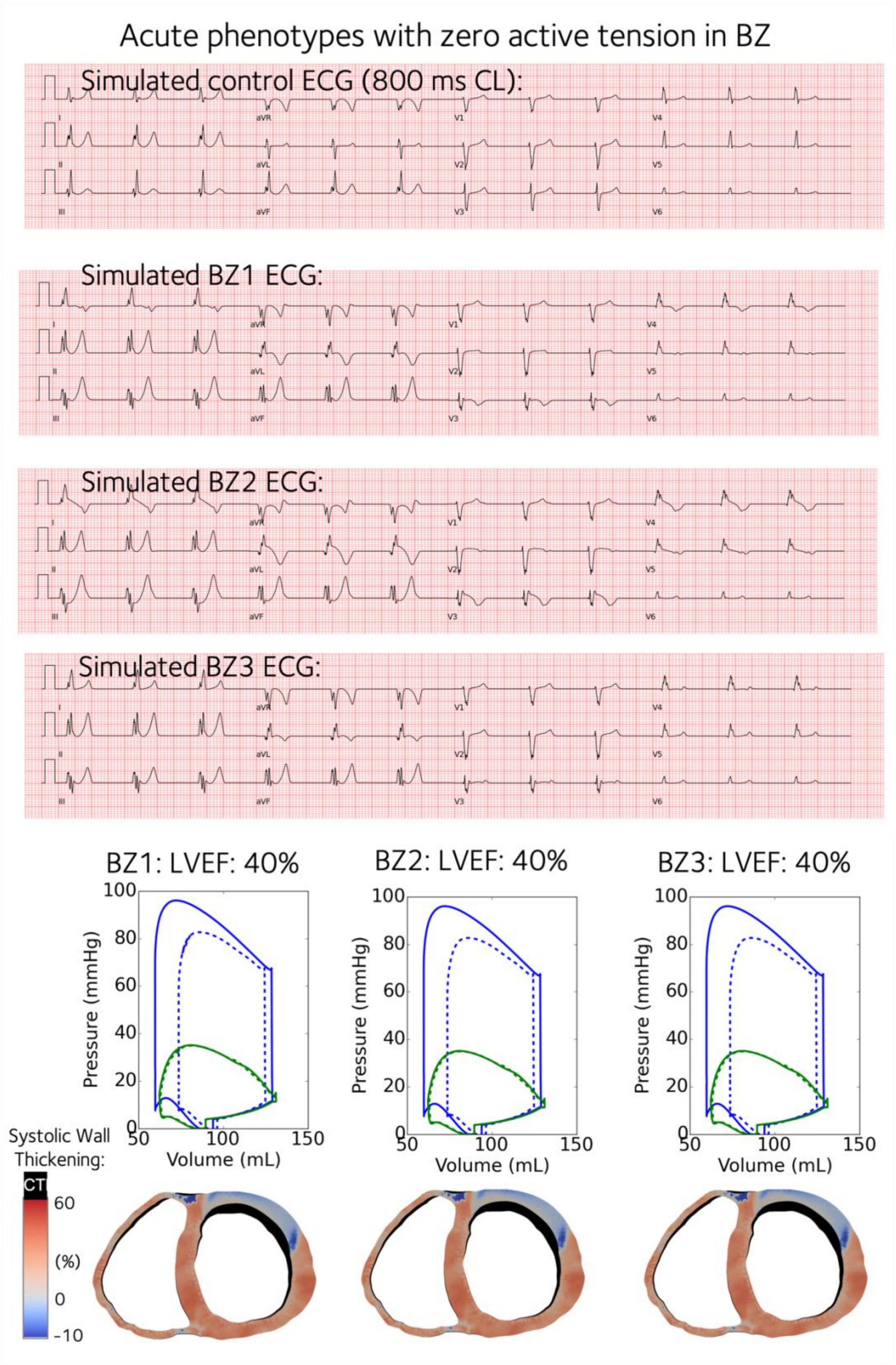


**Figure S9:** Simulated acute stage 12-lead ECGs for Acute BZ1-3 with contractility turned off in the BZs. Acute BZ1 caused T wave inversion in precordial leads of V3 and V4, where the QT prolongation was more significant. Acute BZ2 caused Brugada phenocopy in leads V3-V5, while Acute BZ3 produced similar ECG morphology as the control case.

**Table S8:** Simulated ECG biomarkers from biventricular electromechanical simulations for the acute and the chronic post-MI stages. For the acute stage, Acute BZ1 and BZ2 caused significant QT prolongation, longer T peak to T end, whereas the Acute BZ3 induced milder effects. For the chronic stage, both Chronic RZ1 and RZ2 led to QT prolongation, with RZ2 also generating longer T wave duration, T peak to T end, and T start to T peak than control and RZ1 at CL=800ms. At fast pacing of CL=500ms, both Chronic RZ1 and RZ2 caused longer QT, T wave and T start to T peak durations. For both stages, the QT dispersions did not reflect the repolarization dispersion very well.

| **ECG biomarkers** | **Control** | **Acute BZ1** | **Acute BZ2** | **Acute BZ3** | **Chronic RZ1** | **Chronic RZ2** | **Control CL=500ms** | **Chronic RZ1 CL=500ms** | **Chronic RZ2 CL=500ms** |
|---|---|---|---|---|---|---|---|---|---|
| QRS duration (ms) | 79±2 | 91±5 | 95±9 | 92±6 | 94±6 | 93±5 | 86±7 | 86±7 | 86±7 |
| T duration (ms) | 96±17 | 163±40 | - | 113±18 | 122±42 | 291±20 | 97±19 | 100±15 | 140±37 |
| T peak to T end (ms) | 59±10 | 101±31 | 122±46 | 68±8 | 68±11 | 153±30 | 57±8 | 55±12 | 82±17 |
| T start to T peak (ms) | 38±8 | 61±30 | - | 45±10 | 54±32 | 138±12 | 40±11 | 45±6 | 58±22 |
| QT interval (ms) | 322±1 | 356±24 | 371±3 | 336±5 | 385±4 | 578±3 | 305±2 | 344±4 | 419±4 |
| QT dispersion (precordial) (ms) | 3 | 55 | 9 | 7 | 7 | 7 | 4 | 6 | 5 |

#### Electrophysiological characteristics of post-MI in more detail

The simulated transmural gradients of activation times, repolarisation times, and action potential duration (APD90) are shown in more detail below. Despite the inclusion of the mid-myocardial cell type that has elevated APD90 in the biventricular model, the transmural profile of APD90 in the simulated APD90 map is monotonically decreasing from endocardium to epicardium due to electronic coupling effects.

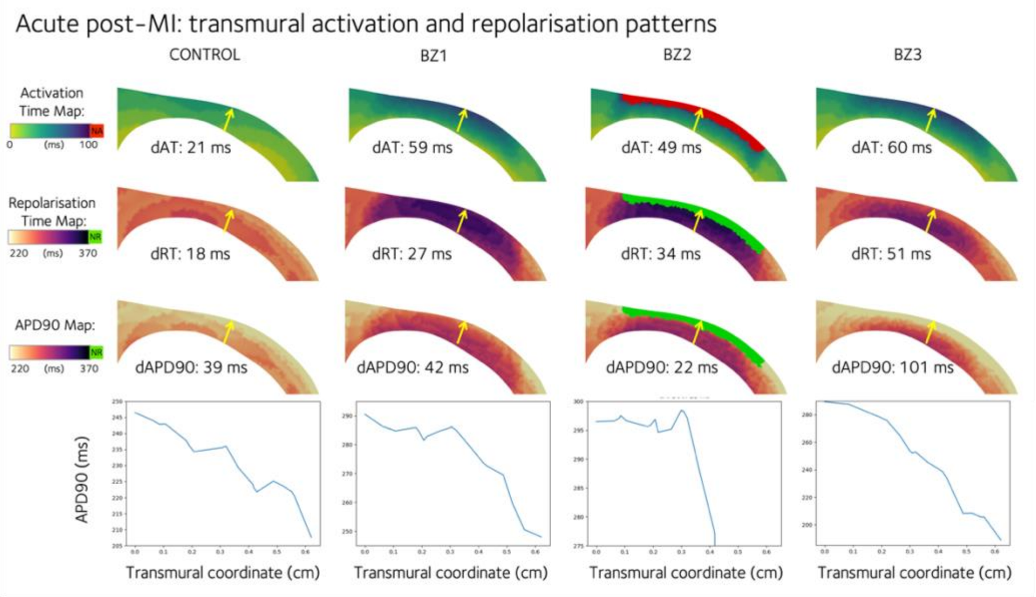


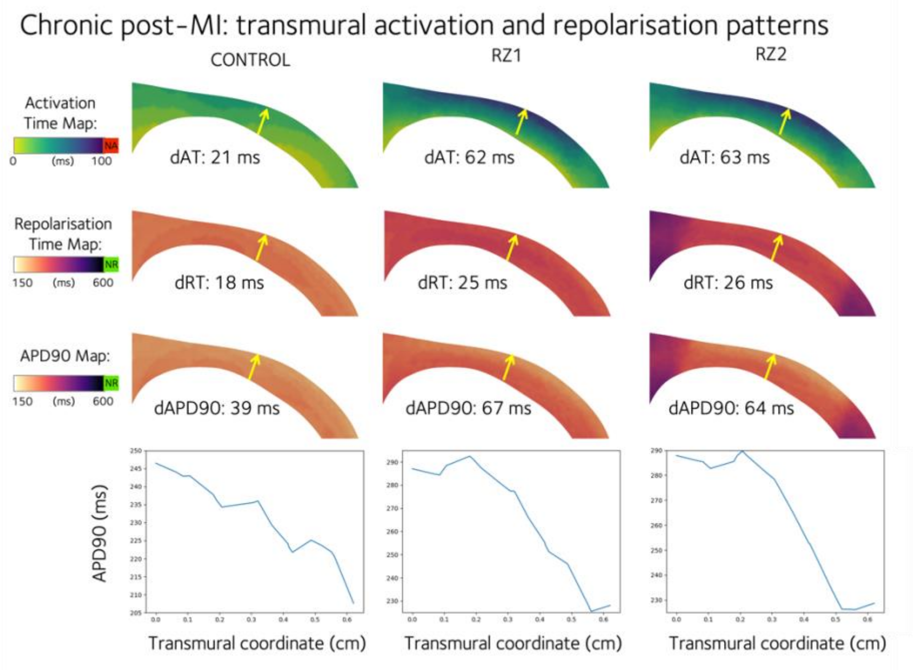


**Figure S10:** Transmural activation time (AT), repolarisation time (RT), and action potential duration at 90% repolarisation (APD90) are shown for the acute (top panel) and chronic (bottom panel) phenotypes. A mid-ventricular anterior transmural slice is taken from the left ventricle that shows the cross-section of the anterior infarction and border zones. The transmural gradient is evaluated as the quantity of interest at the epicardium minus that at the endocardium, and is given as dAT, dRT, and dAPD90 values for each cross-section, evaluated at the beginning and end of the yellow arrows. APD90 is plotted across a transmural line as indicated by the yellow arrow.

#### Explanation of the BZ2 activation pattern

Compared to the other cases, the acute BZ2 had the strongest inhibition of the L-type calcium current, and therefore the least safe epicardial conduction. Due to the transmural differences of the L-type calcium channel expression, the infarct zone in the midmyocardium had a higher level of the calcium current than the epicardial BZ, which explained the more robust conduction in the infarct zone than in the BZ.

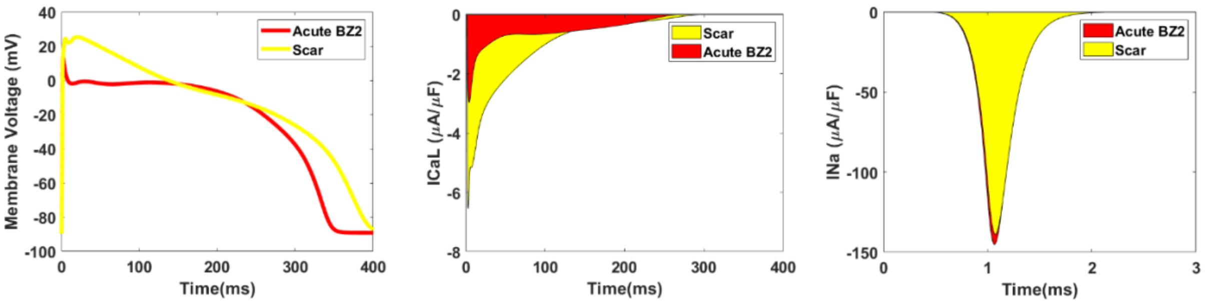


**Figure S11:** Single cell simulations of the action potential (left), L-type calcium ionic current (middle) and sodium ionic current (right) compared between the scar region and the acute BZ2 help to explain the activation pattern in BZ2.

**Table S9:** Simulated pressure-volume mechanical biomarkers for each heart beat from biventricular electromechanical simulations for acute and chronic stages post-MI: left and right end diastolic volumes (EDVL, EDVR), left and right stroke volumes (SVL, SVR), left and right ventricular ejection fractions (LVEF, RVEF). For the acute stage, Acute BZ1 and Acute BZ3 generated the same degree of reduction in SVL and LVEF, whereas the Acute BZ2 induced the smallest SVL and LVEF. For the chronic stage, both Chronic RZ1 and Chronic RZ2 produced the same SVL and LVEF at both pacing rates despite their difference in the degree of repolarization heterogeneity.

| | **Pressure-volume Biomarkers** | **Control** | **Acute BZ1** | **Acute BZ2** | **Acute BZ3** | **Chronic RZ1** | **Chronic RZ2** |
|---|---|---|---|---|---|---|---|
| 800 ms CL | EDVL (mL) | 129, 129, 129 | 124,124,124 | 124,125,125 | 124,124,124 | 127, 126, 126 | 127, 126, 126 |
| | EDVR (mL) | 130,131, 131 | 130,131,131 | 130,131,131 | 130,131,131 | 132,133,133 | 132,133,133 |
| | SVL (mL) | 68, 69, 69 | 59,59,59 | 53,53,53 | 59,59,59 | 62, 61, 61 | 62, 61, 61 |
| | SVR (mL) | 68,68,68 | 67,67,67 | 67,67,67 | 67,67,67 | 69,69,69 | 69,69,69 |
| | LVEF (%) | 53,53,53 | 47,47,47 | 43,43,43 | 47,47,47 | 49, 48, 48 | 49, 48, 48 |
| | RVEF (%) | 52,52,52 | 51,51,51 | 51,51,51 | 51,51,51 | 52,52,52 | 52,52,52 |
| | Peak left systolic pressure (kPa) | 12,12,12 | 11,11,11 | 11,11,11 | 11,11,11 | 11, 11, 11 | 11, 11, 11 |
| | Peak right systolic pressure (kPa) | 4,4,4 | 4,4,4 | 4,4,4 | 4,4,4 | 4,4,4 | 4,4,4 |

| | | | | | |
|---|---|---|---|---|---|
| 500 ms CL | EDVL (mL) | 111,112, 113, 113,113, 113 | NA | 114,107,111, 108,110,108 | 116,108,111, 109,110,109 |
| | EDVR (mL) | 123,122, 123, 123,123, 123 | | 125,117,122, 119,121,119 | 126,119,123, 120,121,120 |
| | SVL (mL) | 53,53,54, 54,54,54 | | 50,43,47, 44,45,44 | 51,43,46, 44,45,44 |
| | SVR (mL) | 61,61,61, 61,61,61 | | 64,56,62, 56,60,57 | 65,57,62, 59,60,59 |
| | LVEF (%) | 47,47,47, 47,47,47 | | 44,39,42, 40,41,40 | 44,39,41, 40,41,40 |
| | RVEF (%) | 49,49,49, 49,49,49 | | 51,47,50, 47,50,48 | 51,48,50, 49,49,49 |
| | Peak left systolic pressure (kPa) | 11,11,11, 11,11,11 | | 10,10,10, 10,10,10 | 10,10,10, 10,10,10 |
| | Peak right systolic pressure (kPa) | 4,4,4, 4,4,4 | | 4,4,4, 4,4,4 | 4,4,4, 4,4,4 |

#### SM7: Post-MI SERCA and CaMKII remodeling promote alternans generations in models with more preserved calcium magnitudes

In our cellular population of post-MI models, higher inducibility of alternans were observed at both the acute and the chronic stages at fast pacing (Figure S12).

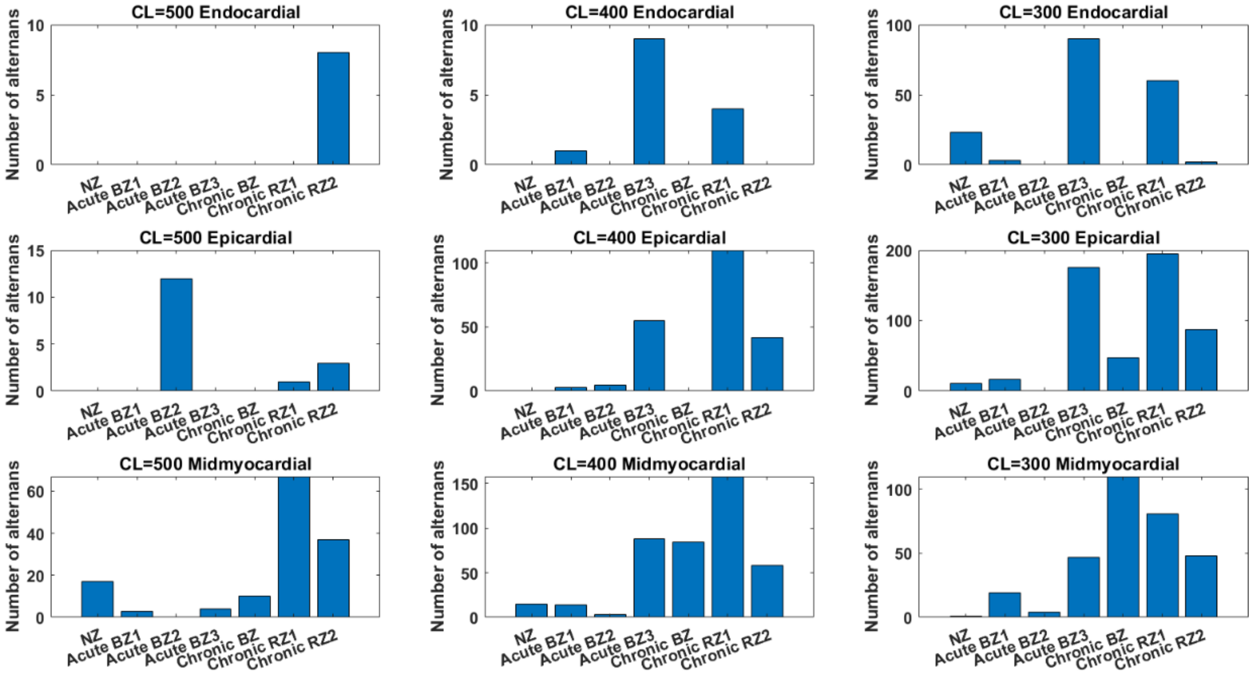


**Figure S12:** The effects of the BZ and RZ remodelling of the acute and chronic stages on alternans generation in the population of 245 population of models at CL=500ms, 400ms and 300ms.

All three types of chronic post-MI remodeling promoted alternans generation, especially in the epicardial and midmyocardial populations (Table S10). Alternans in the midmyocardial layer were mostly due to EADs, whereas the epicardial alternans were repolarization alternans (Figure S13 and S14).

**Table S10:** Number of alternans induced by three chronic remodelling at CL=500, 400 and 300 ms in endocardial, midmyocardial and epicardial population of models.

| Population of models (n=245) | No. of alternans at CL=300ms | No. of alternans at CL=400ms | No. of alternans at CL=500ms | Key parameters for alternans |
|---|---|---|---|---|
| Chronic BZ | Mid (110) > Epi (47) | Mid only (84) | Mid only (10) | $\uparrow G_{CaL}$, $\uparrow G_{Kr}$, $\uparrow P_{Jup}$ |
| Chronic RZ1 | Epi (195) > Mid (81) > Endo (60) | Mid (158) > Epi (110) > Endo (4) | Mid (67) > Epi (1) | $\uparrow G_{CaL}$, $\uparrow P_{Jup}$ |

| | | | | |
|---|---|---|---|---|
| Chronic RZ2 | Epi (88) > Mid (48) > Endo (2) | Mid (58) > Epi (42) | Mid (37) > Endo (8) > Epi (3) | ↑$G_{CaL}$, ↑$G_{Kr}$, ↓$G_{NCX}$, ↑$P_{Jup}$, ↑$P_{Jrel}$ |

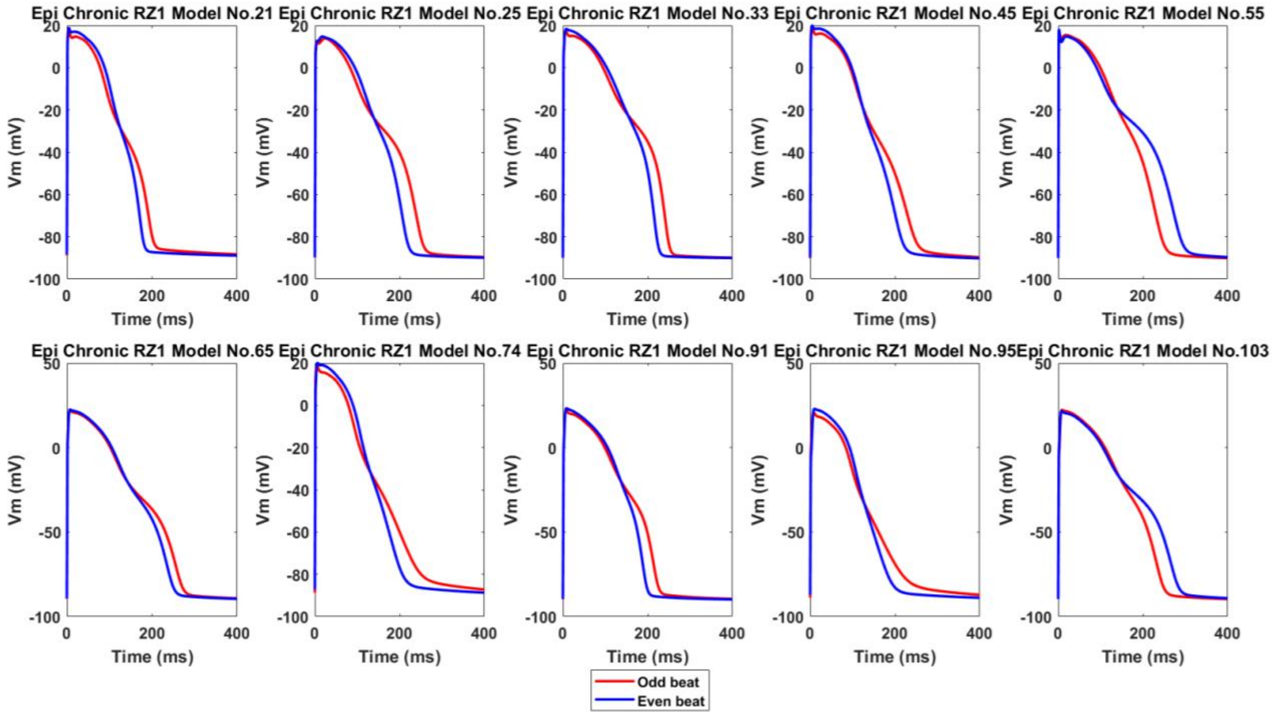


**Figure S13:** Ten representative alternans in the epicardial population of Chronic RZ1, showing calcium-driven repolarization alternans without EADs.

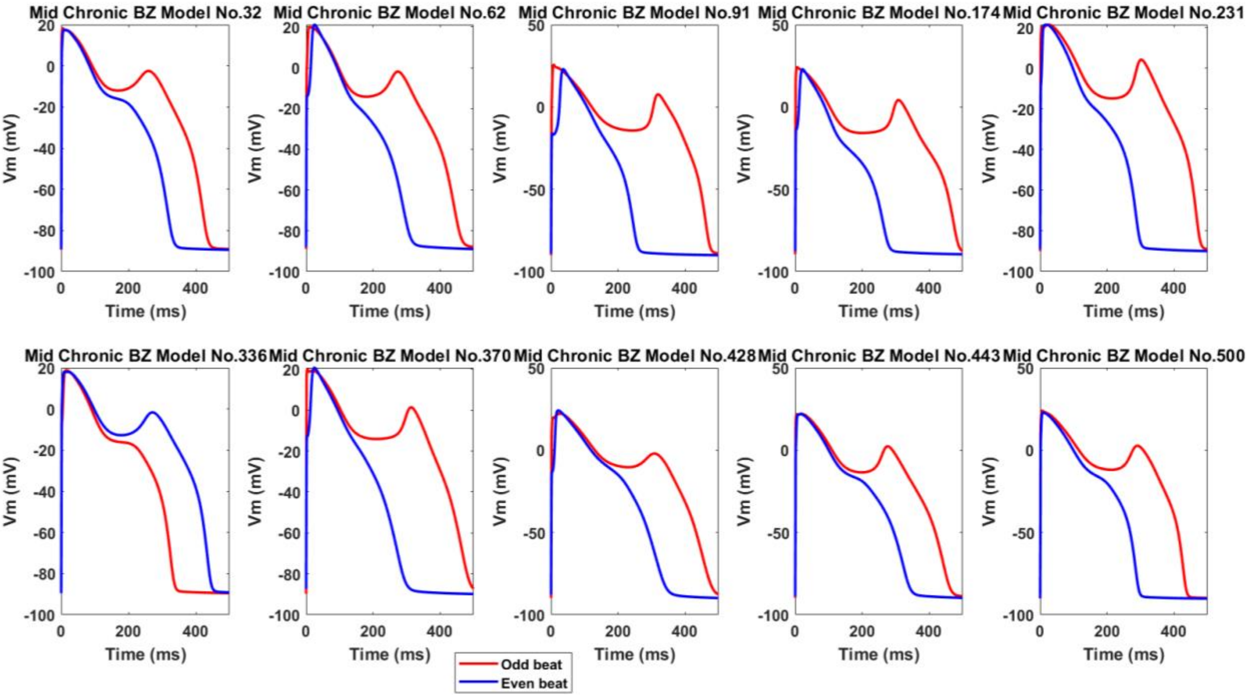


**Figure S14:** Ten representative alternans in the midmyocardial population of Chronic BZ, showing EAD as a major cause of big alternans.

Here we investigated: 1) what are the key individual post-MI ionic current remodeling contributing to the generation of alternans, and 2) what are the underlying ionic currents in the population of models that determine whether a post-MI model is prone for alternans induction?

In order to illustrate the underlying mechanisms, the baseline chronic remote zone model (with Chronic RZ1 remodeling) was chosen as an example in Figure S15. At CL=300 ms, the model had alternations of long and short APDs in the odd and even beats (blue solid traces), and it was clear that the alternans was associated with the insufficient calcium re-uptake and slow calcium recovery in junctional sarcoplasmic reticulum (JSR). When the inhibition of SERCA pump ($J_{up}$) was switched off (red dashed traces), alternans disappeared along with a significant increase of the JSR calcium level. Apart from the insufficient calcium re-uptake in the remodeling, higher CaMKII activation and slower calcium release further contributed to the alternans. When CaMKII activation and $J_{rel}$ kinetics were switched back to normal (yellow dashed traces), the duration of calcium release was shorter, and there was more time for JSR calcium to recover before the next beat, leading to the elimination of alternans (yellow dashed traces).

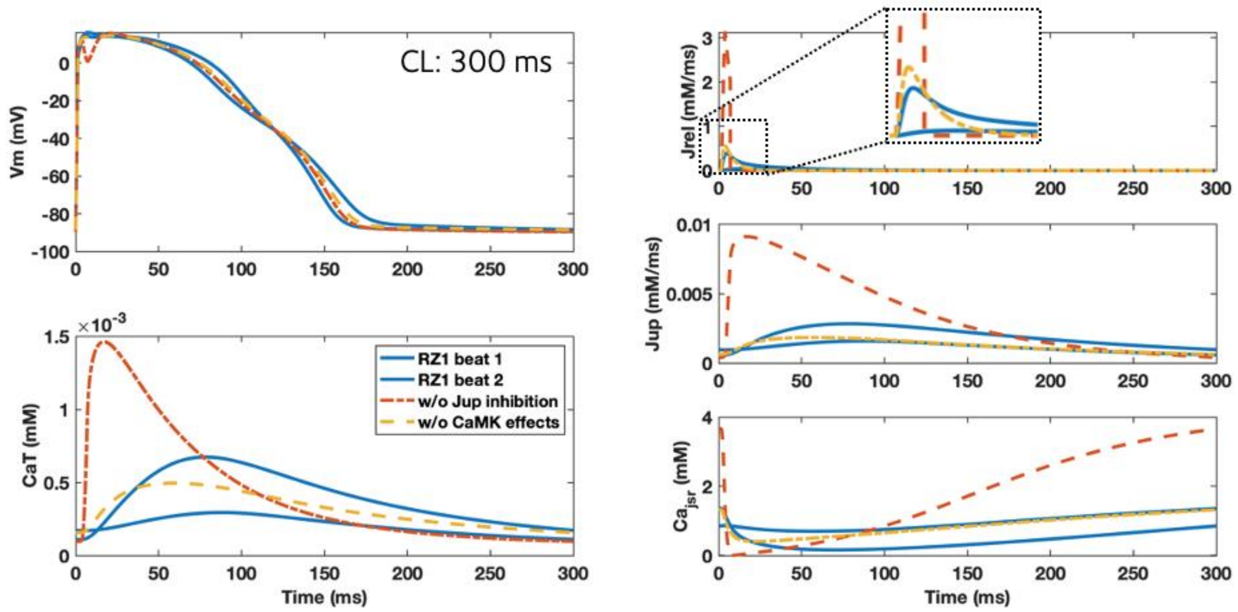


**Figure S15:** Inhibition of $J_{up}$ and slower calcium release ($J_{rel}$) caused by enhanced CaMKII activity promoted alternans.

On the other hand, $I_{KCa}$ augmentation played a protective role against alternans generation by shortening APD and therefore regulating calcium dynamics (Figure S16, S17):

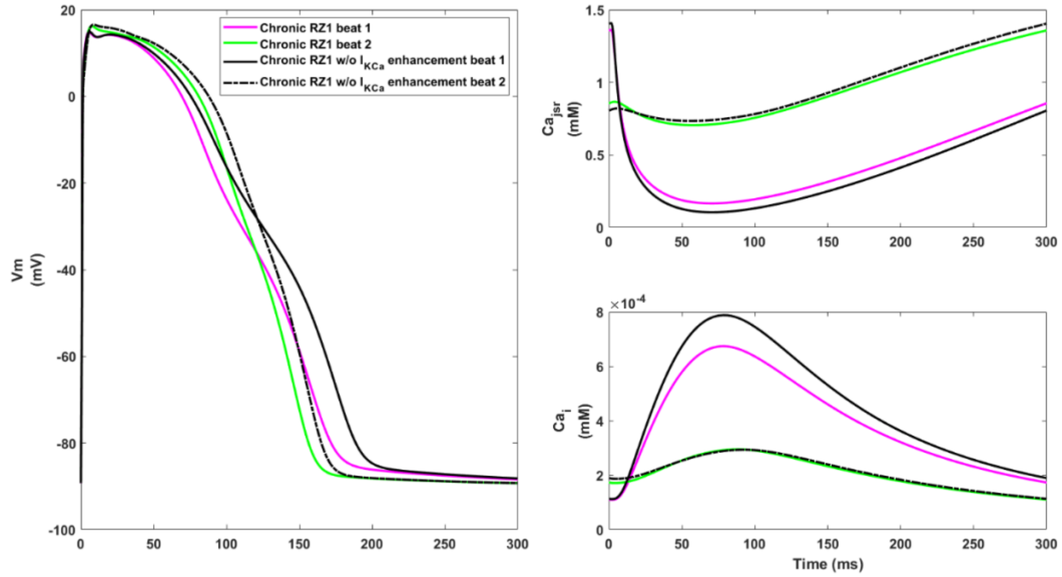


**Figure S16:** Effects of $I_{KCa}$ enhancement on alternans generation in the chronic stage. Left: switching $I_{KCa}$ activity back to normal (black traces) caused AP prolongation and bigger alternans. Weaker $I_{KCa}$ also led to stronger CaT (bottom right, black solid line) and larger calcium release in the longer beat (black solid line) that was more difficult for calcium level to recover in JSR (upper right, black solid line).

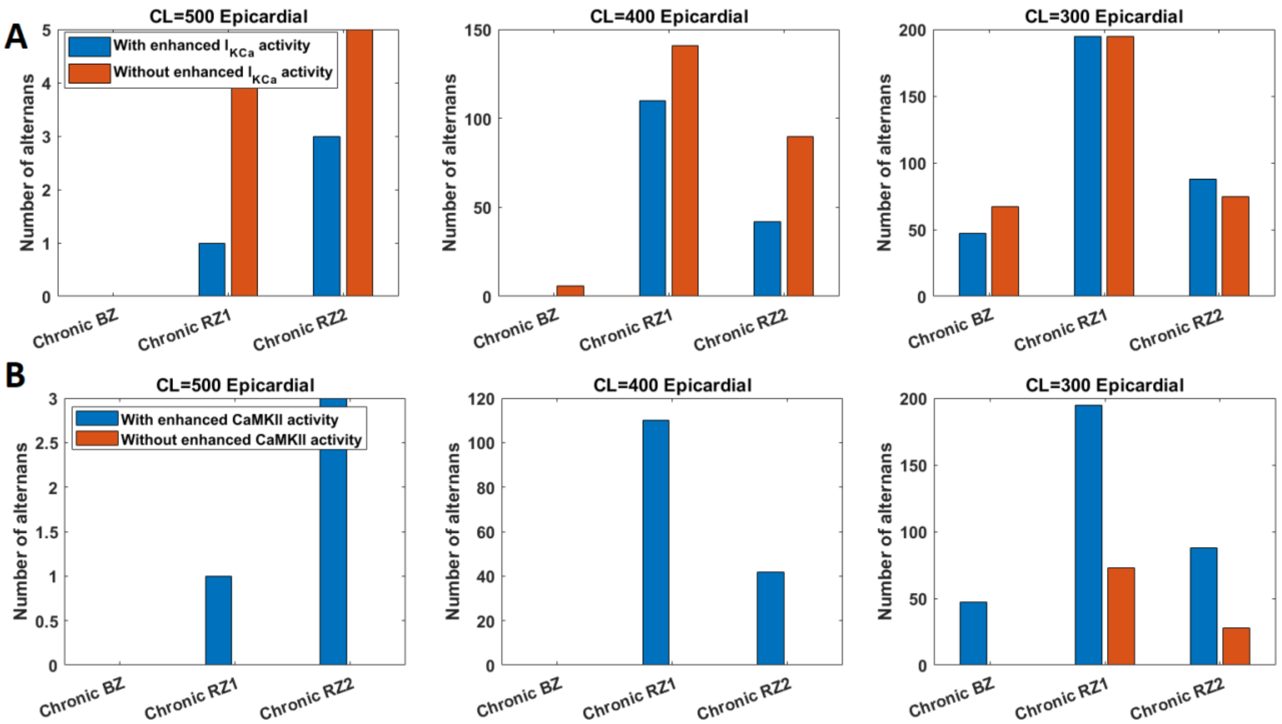


**Figure S17:** CaMKII and $I_{KCa}$ had opposite roles on alternans inducibility in chronic ionic remodelling. Enhanced $I_{KCa}$ tended to inhibit alternans generation (A), whereas augmented CaMKII promoted alternans (B).

As stronger $G_{CaL}$ and stronger $P_{Jup}$ were consistently observed in the chronic population of alternans models (Table S10), the effects of reversing these trends were investigated. In a representative alternans model, when $G_{CaL}$ was inhibited by 20% (Figure S18, the purple trace), the smaller calcium influx led to weaker calcium release, causing a milder reduction of JSR calcium level that was easier to refill, and leading to the elimination of alternans. If $J_{up}$ was further inhibited by 20% (Figure S18, the green trace), the slower calcium re-uptake led to lower initial JSR calcium level at the beginning of a beat, resulting in a smaller $J_{rel}$ and a milder JSR calcium reduction that was also easier to refill. However, $G_{CaL}$ and $J_{up}$ inhibition suppressed alternans generation at the cost of reducing CaT magnitude (Figure S18, CaT panel).

Although the chronic remodeling in the RZ decreased CaT amplitude compared with the NZ, the alternans models among the remodeling population can have relatively preserved CaT amplitudes (Figure S19). The relatively higher $G_{CaL}$ and $P_{Jup}$ in the alternans models contribute to bigger $CaT_{max}$ and smaller $CaT_{min}$ (Table S10, Figure S19). This suggests that models that were prone to alternans development may display a relative preserved CaT amplitude and LVEF.

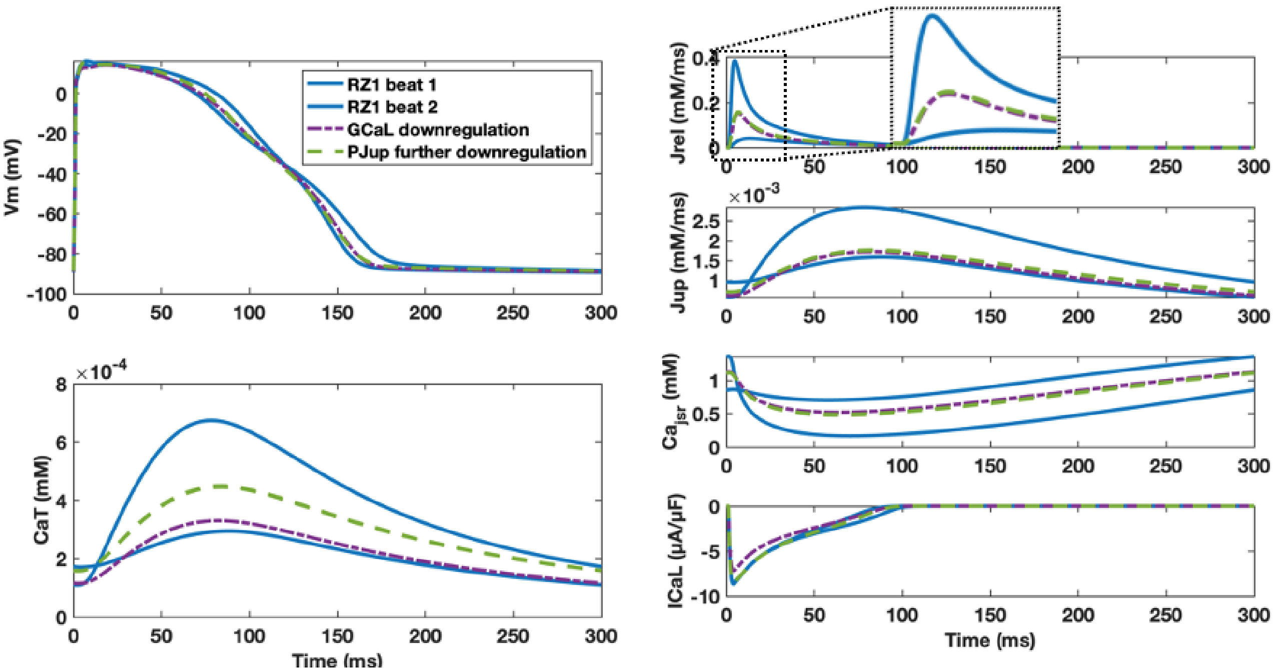


**Figure S18:** With chronic post-MI ionic remodelling, alternans models needed stronger $G_{CaL}$ and more preserved $P_{Jup}$ to enable alternans generation.

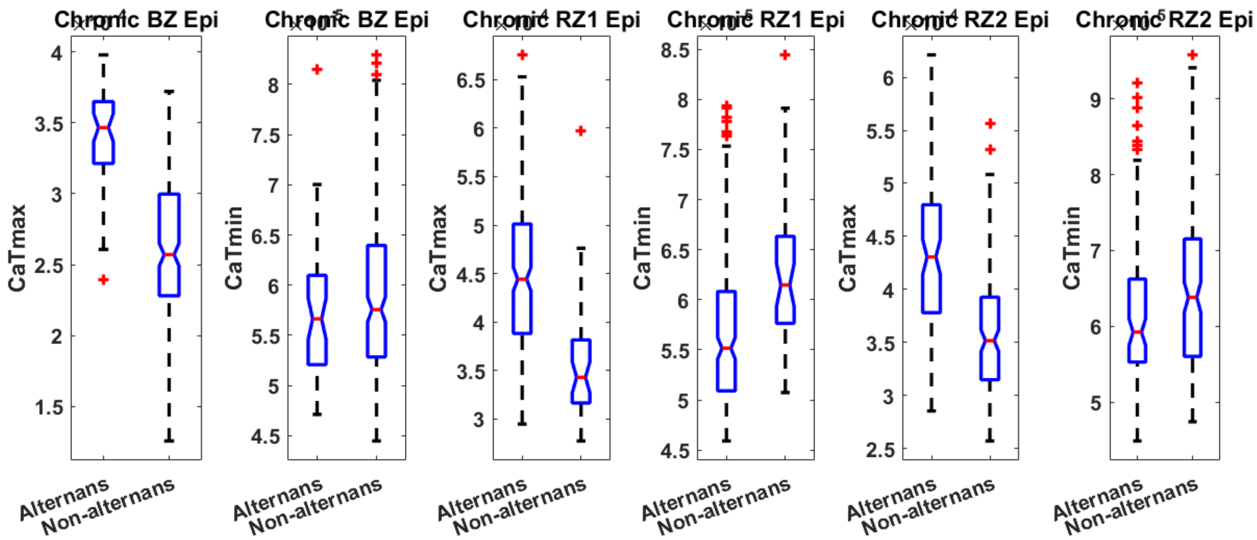


**Figure S19:** Alternans models had bigger $CaT_{max}$ than the non-alternating models in the epicardial populations with chronic post-MI remodelling (all with $p<0.001$). In addition, alternans models tended to have smaller $CaT_{min}$ in epicardial populations of Chronic RZ1 ($p<0.001$) and RZ2 ($p<0.05$), while the difference was not statistically significant for Chronic BZ.

#### SM8: Post-MI $I_{Kr}$ and $I_{NaL}$ remodeling promote EAD generations in models with preserved calcium magnitudes

All three types of chronic post-MI ionic remodeling promoted EADs and repolarization failure (RFs), and midmyocardium was most prone for the development of EAD (Figure S20, Table S11).

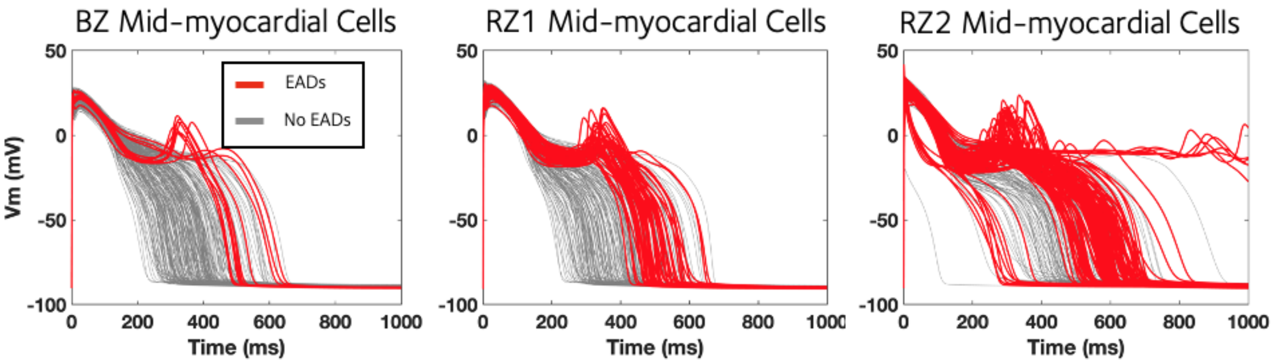


**Figure S20:** Midmyocardium was most prone for the development of EAD under chronic post-MI remodelling.

Cellular early afterdepolarizations (EADs) or repolarization failure (RF) were promoted in the post-MI population of chronic models, and we investigated the following two questions: 1) what are the key individual post-MI ionic remodeling contributing to the generation of EADs, and 2) what are the underlying ionic currents that determine whether a post-MI model is prone to EAD development?

To illustrate the effects of the individual chronic ionic remodeling on the inducibility of EADs, a representative model was chosen from the population (Figure S21), which had a normal AP at a CL of 1000 ms in NZ (the blue trace). When the Chronic BZ ionic remodeling was introduced, an EAD was generated (the red trace). Removing the $I_{NaL}$ remodeling did not eliminate the EAD (the yellow trace), and similarly when $I_{Kr}$ inhibition was removed, EAD was still maintained (the purple trace). However, when both $I_{NaL}$ augmentation and $I_{Kr}$ inhibition were absent, the EAD was eliminated (the green trace). Although $I_{NaL}$ and $I_{Kr}$ remodeling were the key factors inducing EAD generation in the chronic stage, $I_{CaL}$ re-activation was also a necessary mechanism (the light blue trace).

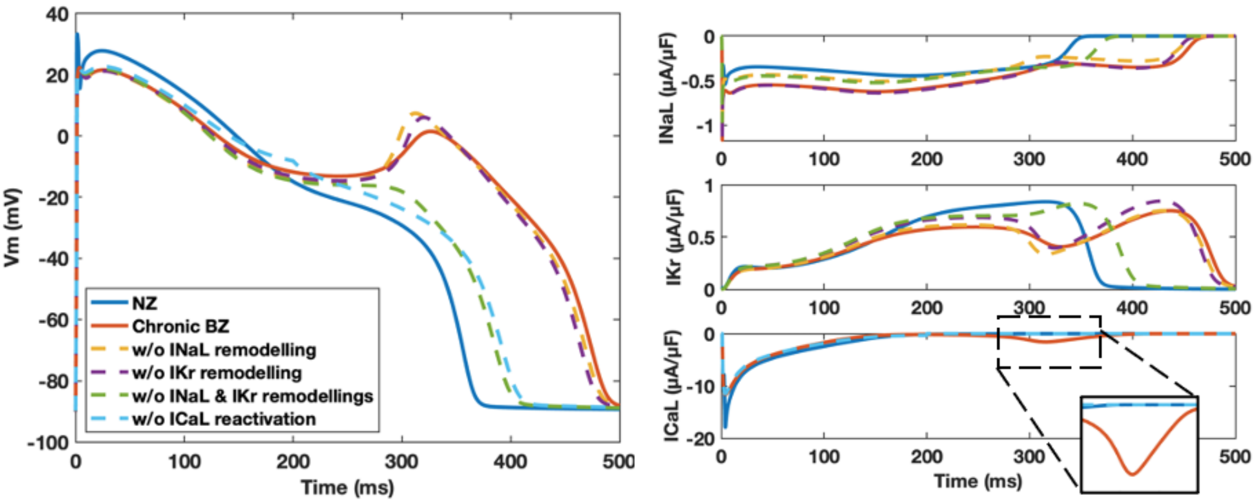


**Figure S21:** Chronic ionic remodelling promotes EAD generation through the enhanced $I_{NaL}$ and suppressed $I_{Kr}$, which facilitate $I_{CaL}$ reactivation.

**Table S11:** Number of EADs and RFs induced by three chronic remodelling in endocardial, midmyocardial and epicardial population of models.

| Population of models (n=245) | No. of EADs and RFs at CL=1000ms | Key parameters for EADs and RFs |
|---|---|---|
| Chronic BZ | Mid only (11) | $\uparrow G_{CaL}$, $\downarrow G_{Kr}$, $\uparrow G_{NCX}$ |
| Chronic RZ1 | Mid only (52) | $\uparrow G_{CaL}$, $\downarrow G_{Kr}$, $\uparrow G_{NCX}$ |
| Chronic RZ2 | Mid (118) > Epi (9) > Endo (1) | $\uparrow G_{CaL}$, $\downarrow G_{Kr}$, $\uparrow G_{NCX}$, $\uparrow P_{Jup}$ |

By comparing the parameters of EAD models against the non-EAD models in the chronic populations, stronger $G_{CaL}$, $G_{NCX}$ and weaker $G_{Kr}$ were consistently observed in the EAD populations (Table S11). Due to the stronger $G_{CaL}$ and $G_{NCX}$, the EAD models also displayed a stronger $CaT_{max}$ and a lower $CaT_{min}$ in all three ionic remodeling populations (Figure S22). Therefore, these results suggested that models which were prone to EAD development may present as a relative preserved LVEF.

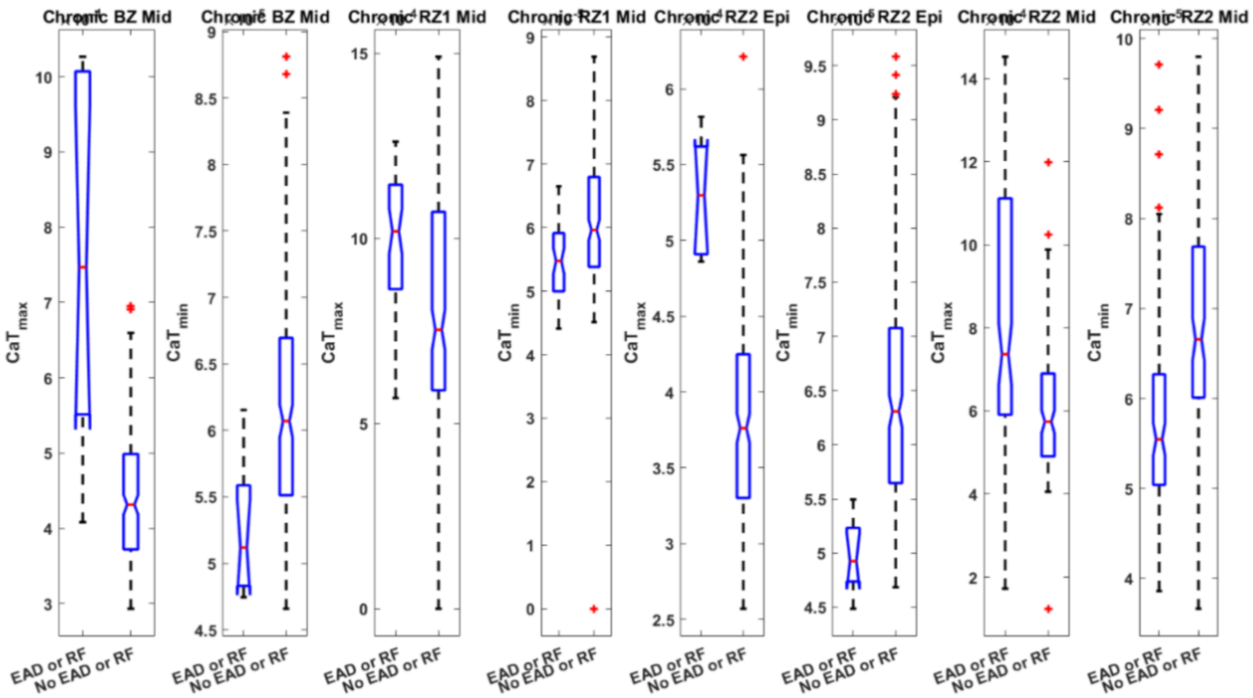


**Figure S22:** EAD models tended to have stronger CaT$_{min}$ and weaker CaT$_{min}$ in the population of Chronic BZ, RZ1 and RZ2 models (all with $p<0.001$).

#### SM9: Both sarcolemmal and calcium dynamics remodeling are necessary for the generation of post-MI EAD alternans

As shown in SM7, EADs can be a major cause of big alternans in the midmyocardial population of chronic post-MI models. Theoretically, calcium alternans can induce EADs under proper conditions, and EADs may display an alternating pattern which occurs in every other beat (Qu & Weiss, 2023). To explore whether the EAD alternans were predominantly EAD or alternans, the 10 representative EAD alternans examples in SM7 were simulated without either EAD related remodelling ($I_{NaL}$ and $I_{Kr}$) or alternans related remodelling ($P_{Jup}$ and CaMKII). EAD alternans disappear when either of the two types of remodelling was switched off (Figure S23-S24), indicating both the sarcolemmal ionic current remodelling and the altered calcium dynamics are necessary conditions for the generation of these EAD alternans.

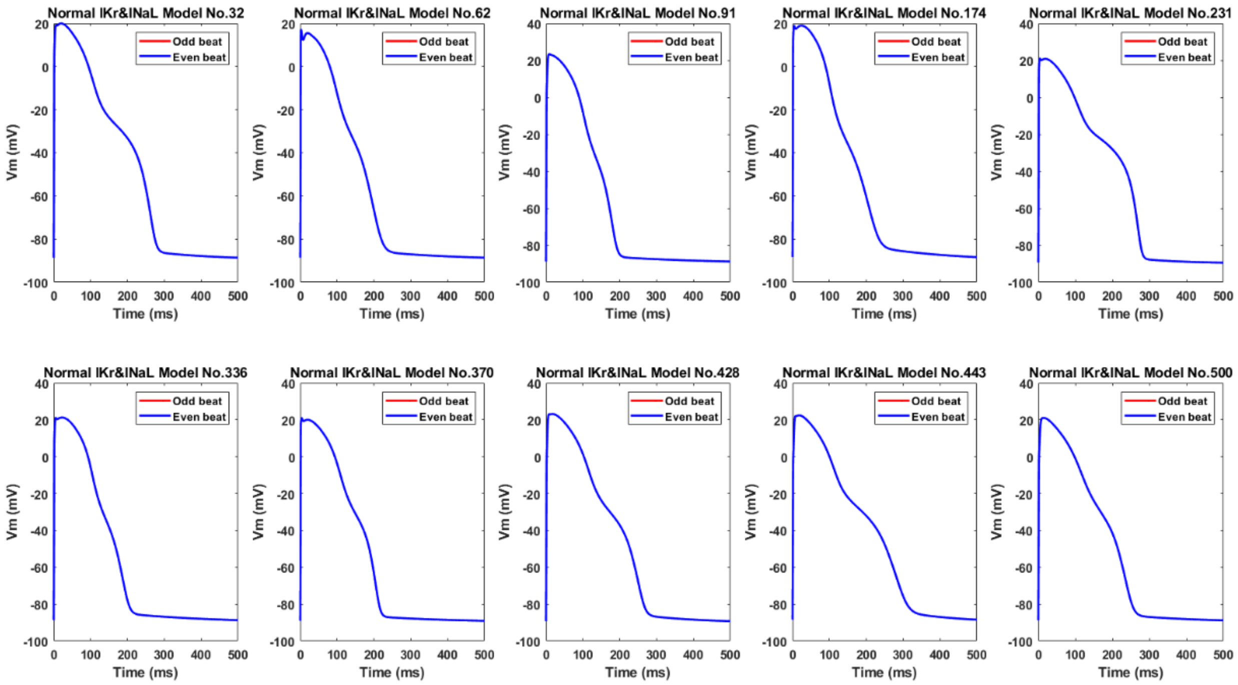


**Figure S23:** Chronic BZ remodelling induced EAD alternans in the ten representative midmyocardial models, but when these models were simulated without $I_{Kr}$ and $I_{NaL}$ remodelling, neither EAD nor alternans occurred.

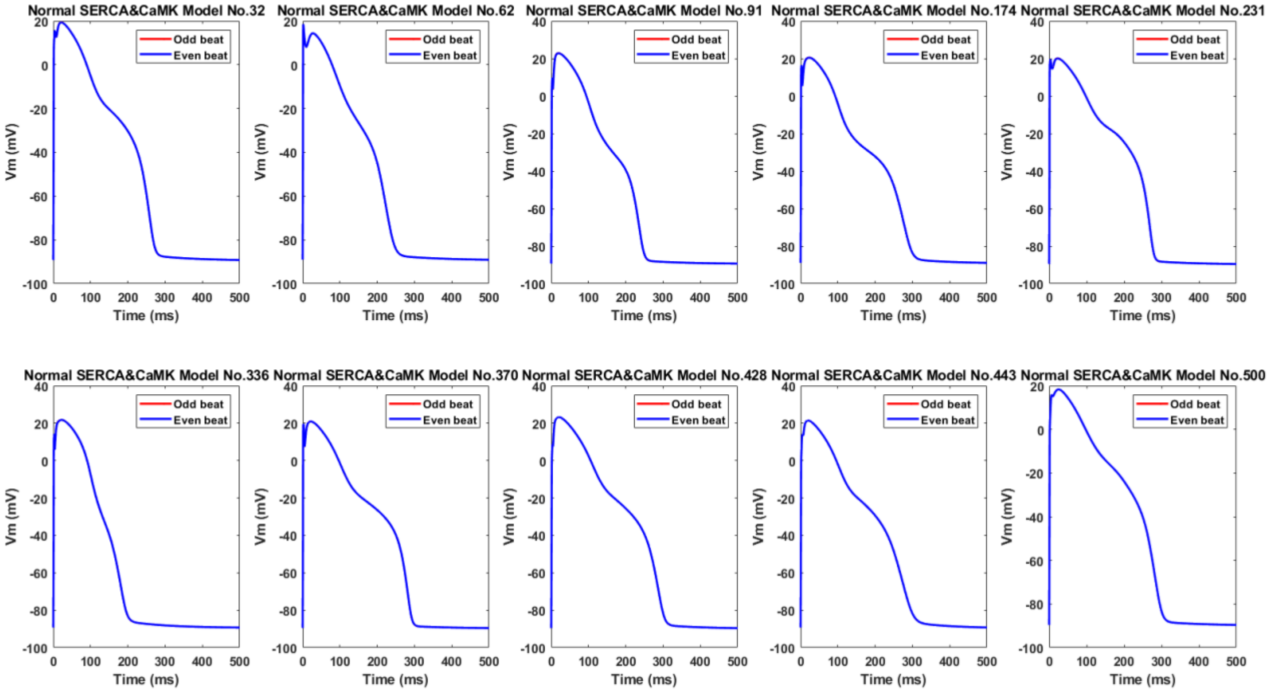


**Figure S24:** Chronic BZ remodelling induced EAD alternans in the ten representative midmyocardial models, but when these models were simulated without $P_{Jup}$ (SERCA) and CaMKII remodelling, neither EAD nor alternans occurred.
